# GRPhIN: Graphlet Characterization of Regulatory and Physical Interaction Networks

**DOI:** 10.1101/2025.02.19.639099

**Authors:** Altaf Barelvi, Oliver Anderson, Anna Ritz

**Author notes:** These authors contributed equally (Altaf Barelvi, Oliver Anderson).

## Abstract

Graphs are powerful tools for modeling and analyzing molecular interaction networks. Graphs typically represent either undirected physical interactions or directed regulatory relationships, which can obscure a particular protein’s functional context. Graphlets can describe local topologies and patterns within graphs, and combining physical and regulatory interactions offer new graphlet configurations that can provide biological insights. We present GRPhIN, a tool for characterizing graphlets and protein roles within graphlets in mixed physical and regulatory interaction networks. We describe the graphlets of mixed networks in *B. subtilis, C. elegans, D. melanogaster, D. rerio*, and *S. cerevisiae* and examine local topologies of proteins and sub- networks related to the oxidative stress response pathway. We found a number of graphlets that were abundant in all species, specific node positions (orbits) within graphlets that were over-represented in stress-associated proteins, and rarely-occurring graphlets that were over-represented in oxidative stress subnetworks. These results showcase the potential for using graphlets in mixed physical and regulatory interaction networks to identify new patterns beyond a single interaction type.

## 1 Introduction

Graphs are powerful tools for computationally representing complex biological networks [8, 15, 37, 22], and are effective in identifying differences between healthy and disease states in organisms [30]. Biological networks can take many forms, and researchers often study networks with isolated interaction types. Some of the most commonly studied network types are protein-protein interaction (PPI) networks and gene regulatory networks (GRNs). PPI networks represent all the physical and molecular interactions of proteins within a system or organism [16] and are represented graphically with proteins as nodes and undirected edges describing their interactions. GRNs differ from PPIs in both the biological context they provide and their graphical structure. GRNs describe interactions between transcription factors (TF) and their target genes and are typically used to study TF-regulated development, diseases, and processes [19].

Many tools represent PPIs and GRNs in isolation, but this can obscure the complete functional context of a particular protein [36]. PPI networks and GRNs do not exist separately: proteins are TFs, genes encode proteins, and physical and regulatory interactions coexist and mix, forming distinct patterns that networks with only one interaction type cannot capture. Wang *et al*. [35] found that physical interactions can participate in regulatory interactions and can even predict long-range enhancer-promoter interactions. Thus, identifying and quantifying patterns and proteins within a heterogeneous network would be valuable in understanding the full biological context of a specific protein or the regulation of a pathway or process.

Characterizing biological networks beyond simple summary statistics has been an ongoing research area. Graphlets are small, connected subnetworks that can describe the local network organization of a graph [25, 24]. Graphlets have been used in many applications of molecular interaction networks [23], including protein function prediction [21], characterizing disease genes [1], and describing the local topologies of signaling pathways [29], among many others.

Since their introduction in the early 2000s, efficient algorithms have been developed to count graphlets and the specific node positions within each graphlet (called *orbits*) [13, 14]. Graphlets have been defined on network extensions such as directed graphs [34, 31, 4], signed graphs [9], and probablistic graphs [11]. While some research describes graphlets for heterogeneous networks, which contain different types of relational information, this area is less well-developed. In 2004, Yeger-Lotem and colleagues counted graphlets that contained a mixture of physical and regulatory interactions in yeast, and identified five graphlets that were over-represented compared to a random rewiring model [36]. In 2017, Sonmez and Can extended graphlets to directed multi-label graphs [33], where edges may have different edge types. In 2020, Dimitrova and colleagues characterized graphlets in undirected multiplex networks with different edge types [10]. To our knowledge, no previous research except for the 2004 work [36] has explicitly considered the case where there are a mix of directed and undirected edges, which is required when combining undirected physical and directed regulatory interactions.

In this paper, we adapt the multi-label graphlet identification work presented in [33] and apply it to mixed networks that contain physical undirected interactions and directed regulatory interactions. Through a new tool called **G**raphlet Characterization of **R**egulatory and **Ph**ysical **I**nteraction **N**etworks (GRPhIN, pronounced “griffin”), we characterize the graphlet distributions of networks of five different species (*B. subtilis, C. elegans, D. rerio, D. melanogaster*, and *S. cerevisiae*), and identify graphlets with a mix of physical and regulatory edges that are common between all species. Additionally, we characterize oxidative stress response, a pathway that is conserved across the five species, both by using the node positions within graphlets as well as by identifying subnetworks that represent the oxidative stress response pathways. GRPhIN gives us new insights into local physical and regulatory patterns within these networks and within the oxidative stress response pathway.

## 2 Methods

We first formally define the graph that we generate from species-specific data. For a particular species, we identify undirected physical relationships (undirected edges) and directed regulatory relationships (directed edges) for a set of proteins (nodes). We build a regulatory and physical interaction (RPI) graph by taking the union of the undirected and directed edges, creating a multigraph (where two nodes may have more than one edge). Here, nodes represent either a protein or the gene that encodes the protein (in the case of a regulatory target). We then relabel the edges in the RPI graph into one of five edge types with seven unique node orbits (Figure 1A), making this graph a multi-label graph with only one edge between any two nodes. Note that this graph still contains a mixture of undirected (e.g., physical-only), bi-directed (e.g., two co-regulating edges), and directed edges. Two RPI graphs are *isomorphic* if there exists an edge-type-preserving bijection. That is, two RPI graphs *A* and *B* with nodes *V*_*A*_ and *V*_*B*_ are isomorphic if there exists a node mapping *f* : *V*_*A*_ *1*→ *V*_*B*_ such that if nodes *u* and *v* are connected by edge type *t* in *A*, then *f* (*u*) and *f* (*v*) are connected by edge type *t* in B (where edge types are shown in Figure 1A). Similarily, an RPI graph *A* is *automorphic* if there is an edge-type-preserving bijection *f* : *A 1*→ *A*.

**Figure 1.**
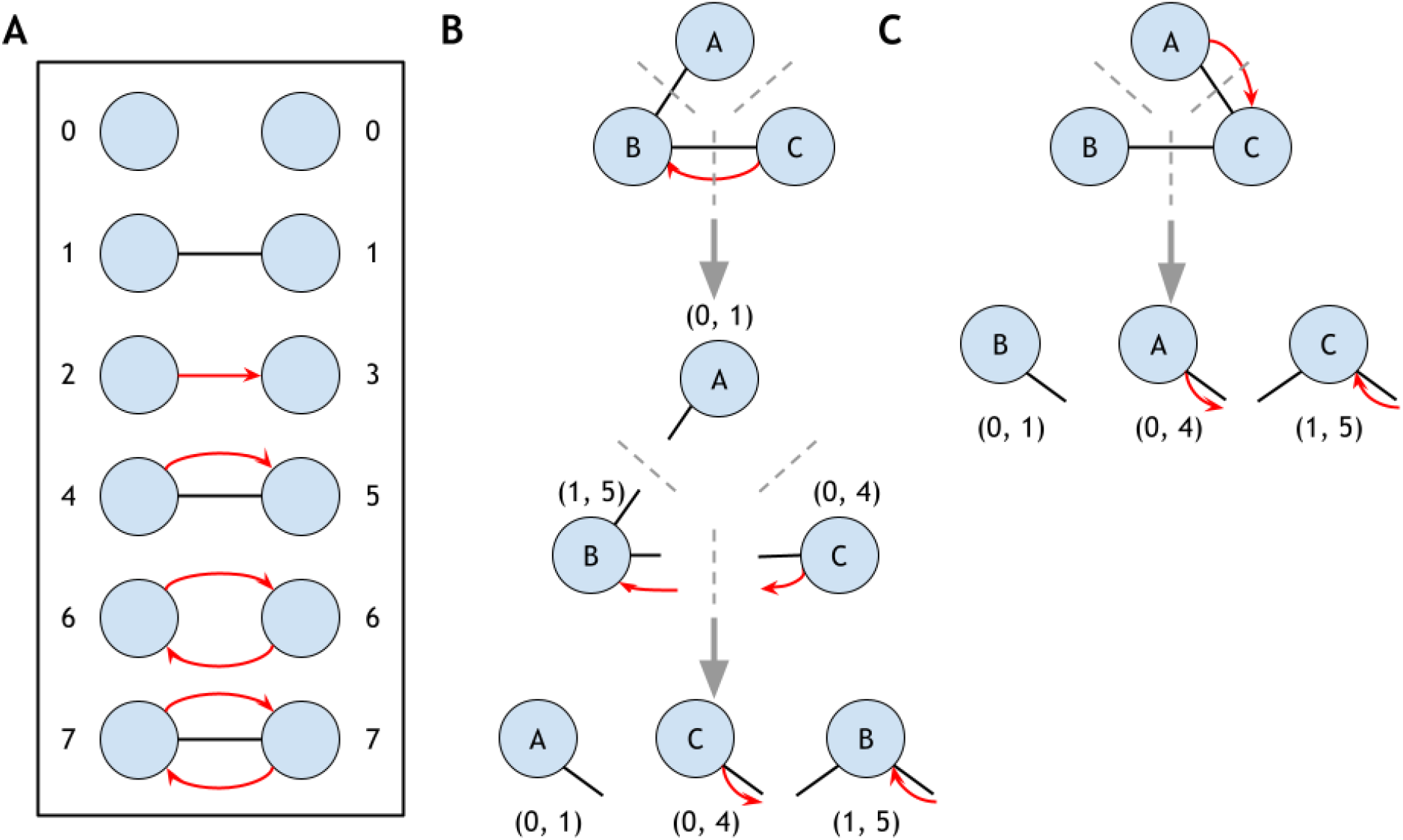
Orbit labeling and graphlet sorting. A) Unique edges and orbit in an RPI graph (orbit 0 indicates a non-edge; orbits 1–7 are mapped to the five distinct edge types). B) Sorting node orbits in ascending order maintains a unique graphlet structure. C) Sorting isomorphisms results in the same unique set of integer pairs.

### 2.1 Regulatory and Physical Interaction (RPI) Graphlets

Given a graph *G*, graphlets are connected, induced, non-isomorphic subgraphs of *G* described in previous work [25, 24, 14]. Graphlets are typically restricted to subgraphs of a specific size (e.g., 2-4 nodes). We define regulatory and physical interaction (RPI) graphlets, which are connected, induced, non-isomorphic subgraphs of the RPI graph. There are five two-node RPI graphlets (corresponding to the five edge types, Figure 1A), but the number of three-node graphlets is much larger than graphlets on directed or undirected graphs: there are 98 three-node graphlets, including 28 that contain two edges (“line graphlets”) and 70 that contain three edges (“triangle graphlets”) (Supplementary Figures S1 and S2). Of these 98 RPI graphlets, 83 (84.7%) contain both physical and regulatory edge types (which we call *mixed graphlets*). We restrict our attention to 2- and 3-node graphlets due to the combinatorial complexity of the enumeration.

For each RPI graphlet, we define the non-automorphic node positions (orbits) which make up that graphlet. Two nodes are equivalent if some automorphism maps one node onto the other; we partition graphlets into equivalence classes defined as orbits [26]. For example, in Figure 1A: orbit 4 reflects a node with a physical edge and outgoing regulatory edge, while orbit 5 reflects a node with a physical edge and incoming regulatory edge. There are seven orbits for the 2-node graphlets and 259 orbits for the 3-node graphlets. While previous work enumerated the orbits, we index the orbits by the specific graphlet for clarity (described in more detail below).

#### 2.1.1 RPI Graphlet Encoding

Since graphlets are connected, each node has exactly one or two incident edges. These incident edges can be described by a node’s orbits in the 2-node RPI graphlets, labeled as index 1–7 (we will call orbits in the 2-node RPI graphlets *edge orbits*). If a node has only one incident edge, we assign a 0 to indicate it is not connected to the other node. Thus, each 3-node RPI graphlet is described as a triple of integer pairs that describe the edge adjacencies among the nodes (Figure 1B/C). For a graphlet *G* with labeled nodes *a, b*, and *c*:

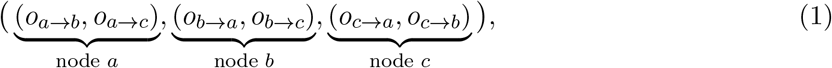

Where

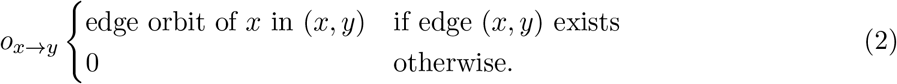

Note that the first two pairs completely determine the graphlet (e.g., once you know the incident edges of *a* and *b*, then *c* is automatically determined), but we keep this notation for ease of exposition. We order the orbits within these pairs: that is, let *o*_*a*_ = sort(*o*_*a*→*b*_, *o*_*a*→*c*_) for graphlet with nodes *a, b*, and *c* (the top portion of Figure 1B illustrates this idea). Without loss of generality, we assume that *a, b, c* are ordered such that *o*_*a*_ *< o*_*b*_ *< o*_*c*_. Thus, the final encoding for graphlet *G* with nodes labeled *a, b, c* is

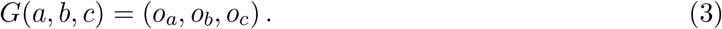

This ordering is unique to each 3-node RPI graphlet and will be used to break symmetries when counting. Figure 1B-C show how two isomorphic graphlets, when sorted, results in the same graphlet encoding.

#### 2.1.2 Counting RPI Graphlets and Orbits

Many analytic and enumeration-based algorithms that are extensions of exact subgraph counting exist to count graphlets (see [26] for a review). However, due to the mixed-graph nature of RPI graphs, we developed GRPhIN to count graphlets. Since we are concerned with 2- and 3-node RPI graphlets, we take an enumeration approach similar to counting directed multi-label graphlets from Sonmez and Can [33].

To count 3-node RPI graphlets, we first arbitrarily order the nodes in the RPI graph *G* and enumerate all connected triples of nodes *i, j, k*. By definition, the subgraph of each connected node triple *i, j, k* is isomorphic to some 3-node RPI graphlet. For each node *i, j*, and *k*, we encode the pairs of node adjacencies and sort each pair to produce *o*_*i*_, *o*_*j*_, *o*_*k*_. Then, we find a mapping *f* : *{i, j, k} 1*→ *{a, b, c}* such that:

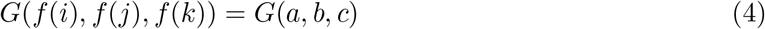

for some graphlet *G* (Figure 1B,C). We increment the count of graphlet *G* for the triple *i, j, k*, and we increment the orbits of *f* (*i*),*f* (*j*), and *f* (*k*) for the nodes *i, j*, and *k*. The output of GRPhIN is (1) a count of every 2-node and 3-node RPI graphlet in the entire RPI graph (called a graphlet distribution vector in previous work [21, 26]) and (2) a count of every orbit in every RPI graphlet for each node in the RPI graph. GRPhIN is available at the https://github.com/Reed-CompBio/grphin.

### 2.2 Algorithm Runtime

We benchmarked enumerating RPI graphlets on the five species-specific networks using three machine configurations: 2020 MacBook Pro with 2 GHz Quad-Core Intel Core i5, 16 GB Memory, and Intel Iris Plus Graphics 1536 MB; 2024 MacBook Pro with Apple M4 chip and 16 GB Memory; GPU machine with AMD Ryzzen Threadripper 3990x 64-Core Processor CPU, 257 GB memory, and two Nvidia GeForm RTX 3060 GPUs (Table 1). *S. cerevisiae* had the longest runtime: 69 minutes (MacBook Pro 2020) and 20 minutes (GPU machine and MacBook Pro 2024). The other networks took an average of 5 minutes to run on a 2020 MacBook Pro, an average of approximately 3.5 minutes on the GPU machine, and the 1.2 minutes on the 2024 MacBook Pro, with the fastest being *B. subtilis* (1.786 seconds). Due to the iterative nature of the algorithm, large graphs with high average node degree will have a long runtime. For these graphs, it is advisable to use a GPU machine to utilize its memory availability, CPU, and GPU power.

**Table 1.** GRPhIN algorithm runtime across five species networks using three machine configurations.

| Organism | Runtime (Seconds) |  |  |
| --- | --- | --- | --- |
|  | MacBook Pro (2020) | MacBook Pro (2024) | GPU Machine |
| *B. sub.* | 6.346 | 1.786 | 3.824 |
| *C. ele.* | 254.277 | 84.817 | 170.845 |
| *D. mela.* | 309.713 | 94.618 | 192.805 |
| *D. rer.* | 381.731 | 110.953 | 229.691 |
| *S. cer.* | 4142.087 | 1240.302 | 1239.425 |

### 2.3 Species-Specific Networks

We collected experimentally, text-mined, and database-validated physical and regulatory interaction data for five non-human model organisms: *B. subtilis, C. elegans, D. melanogaster, D. rerio*, and *S. cerevisiae* (Table 2 and Supplementary Section S1). These interactions come from species-specific databases (e.g., DRoID, FlyBase, WormBase, ZFin, and SubtiWiki) and broader databases that contain species-specific components (e.g., STRING, BIOGRID, and TFLink). These publicly available networks are described in more detail in the ProteinWeaver paper [3].

**Table 2.** Regulatory and physical interaction (RPI) networks. PPI: protein-protein interaction. Reg.: regulatory. Oxidative stress proteins were collected from the Gene Ontology (Methods).

| Organism | Proteins |  | Interactions |  |
| --- | --- | --- | --- | --- |
|  | Total | Stress | PPI | Reg. |
| *B. sub.* | 3,163 | 18 | 6,441 | 5,634 |
| *C. ele.* | 4,106 | 103 | 13,915 | 78,223 |
| *D. mela.* | 12,823 | 145 | 233,054 | 17,530 |
| *D. rer.* | 16,606 | 154 | 45,003 | 25,960 |
| *S. cer.* | 7,644 | 124 | 164,432 | 237,315 |

### 2.4 Oxidative Stress Response Proteins

To identify proteins involved in oxidative stress responses present in all species, we examined gene ontology (GO) annotations [2, 5]. Using AMIGO [7], we exported a list of proteins annotated with the GO term “response to oxidative stress” (GO:0006979), a process common to the five species studied (Table 2). We performed two analyses: a graphlet-based over-representation analysis on oxidative stress subnetworks and an orbit-based over-representation analysis on oxidative stress response proteins.

#### 2.4.1 Graphlet Over-Representation in Oxidative Stress Subnetworks

We wanted to identify over-represented graphlets in oxidative stress response pathways; however, the induced subgraph of oxidative stress proteins on the RPI graphs produced fragmented sub-networks (Supplementary Section S4). Therefore, we used random walks with restarts (RWR) to identify connected subnetworks with a large proportion of oxidative stress response proteins. For each species, we performed a random walk (RWR) on the RPI network restarting at the oxidative stress response proteins with restart probability *α* = 0.85 to generate a list of proteins near oxidative stress response proteins in the graph. We took the induced subnetwork of the top nodes from the RWR, which included 80% of the oxidative stress response proteins (which we will refer to as the *oxidative stress subnetwork*) for each species. We then counted RPI graphlets in each of the oxidative stress subnetworks.

To evaluate whether the frequency of a graphlet in an oxidative stress response network was significant, we performed a network perturbation analysis adapted from Shen-Orr *et al*. [32] and Yeger-Lotem *et al*. [36]. The algorithm allows edge swaps when the four incident nodes meet a “switch condition” that preserves all degress of all edge types (Supplementary Section S5). We generated 1,000 randomized networks with the same node degree and edge counts as the original network for each species, performing enough edge swaps to saturate the number of new edges added (Supplementary Figures S5 and S6). To calculate significance, we compared the counts of each graphlet found in the oxidative stress response networks to the 1,000 randomized networks. If a graphlet was found to be more common compared to the oxidative stress response network in fewer than 10 randomized networks, it was deemed significantly over-represented (*p <* 0.01).

There are other ways to calculate graphlet/motif enrichment, and we also performed an analytical enrichment analysis based on the global networks of each species (Supplementary Section S6). However, we found that this analysis tended to favor sparser graphlets.

#### 2.4.2 Orbit Over-Representation of Oxidative Stress Proteins

We also wanted to identify certain orbits (or roles) within RPI graphs that tend to be occupied by oxidative stress proteins. For this analysis, we performed a random sampling method following a process similar to Agrawal *et al*. [1]. For each orbit in each species, we calculated the observed median orbit count for the oxidative stress proteins and compared that value to the median orbit counts of 1,000 randomly sampled node sets of the same size. An orbit is significantly over-represented in the oxidative stress proteins if less than 1% of the random medians are equal to or larger than the observed median, corresponding to a *p*-value of 0.01.

To further investigate the biological roles of oxidative stress-related proteins within network structures, we examined the biological processes associated with oxidative stress proteins occupying specific orbits. For each significant orbit, we compiled a set of genes and proteins that occupy the orbit and performed Gene Ontology (GO) enrichment analysis [20]. Over-represented GO terms with a false discovery rate (FDR) *<* 0.01 were retained. Given the high number of orbits (259 total) for each species, we narrowed our scope to significant orbits that are part of mixed graphlets.

## 3 Results

### 3.1 RPI Graphlet Characterization of Species Networks

We first examined the distribution of graphlet counts across the five species networks. Of the two-node graphlets (the edge types illustrated in Figure 1A), the edge type that denotes a single physical interaction and the edge type that denotes a single regulatory interaction dominate the RPI graphlet counts: only 0.03-1.1% of the species networks contain the two mixed edge types. These rare edge types appear in some significant RPI graphlets, however, and many mixed graphlets consist of single-physical or single-regulatory edges, as we will see.

Despite the differences in the RPI graph sizes and total abundance of RPI graphlets in each graph, there is a distinct pattern of three-node RPI graphlets across species (Figure 2A). The most abundant RPI graphlets across all species are a line of two physical edges, *G*_1_, and a line with two outgoing regulatory edges from the central node, *G*_24_ (Figure 2B and Supplementary Figure S3). Interestingly, *G*_16_, which includes a mixed physical and regulatory edge, is also abundant in all species (except *D. melanogaster*, where it is present, but not one of the most abundant graphlets), despite mixed graphlets making up a small proportion of the total enumerated graphlets (Supplementary Table S1). Although less abundant compared to line graphlets (Supplementary Table S2), several triangle graphlets are commonly found in all networks (Figure 2B). Note that there is some dependence between the line and triangle graphlets (e.g., many of the triangle graphlets contain subgraphs that are isomorphic to the most common line graphlets).

**Figure 2.**
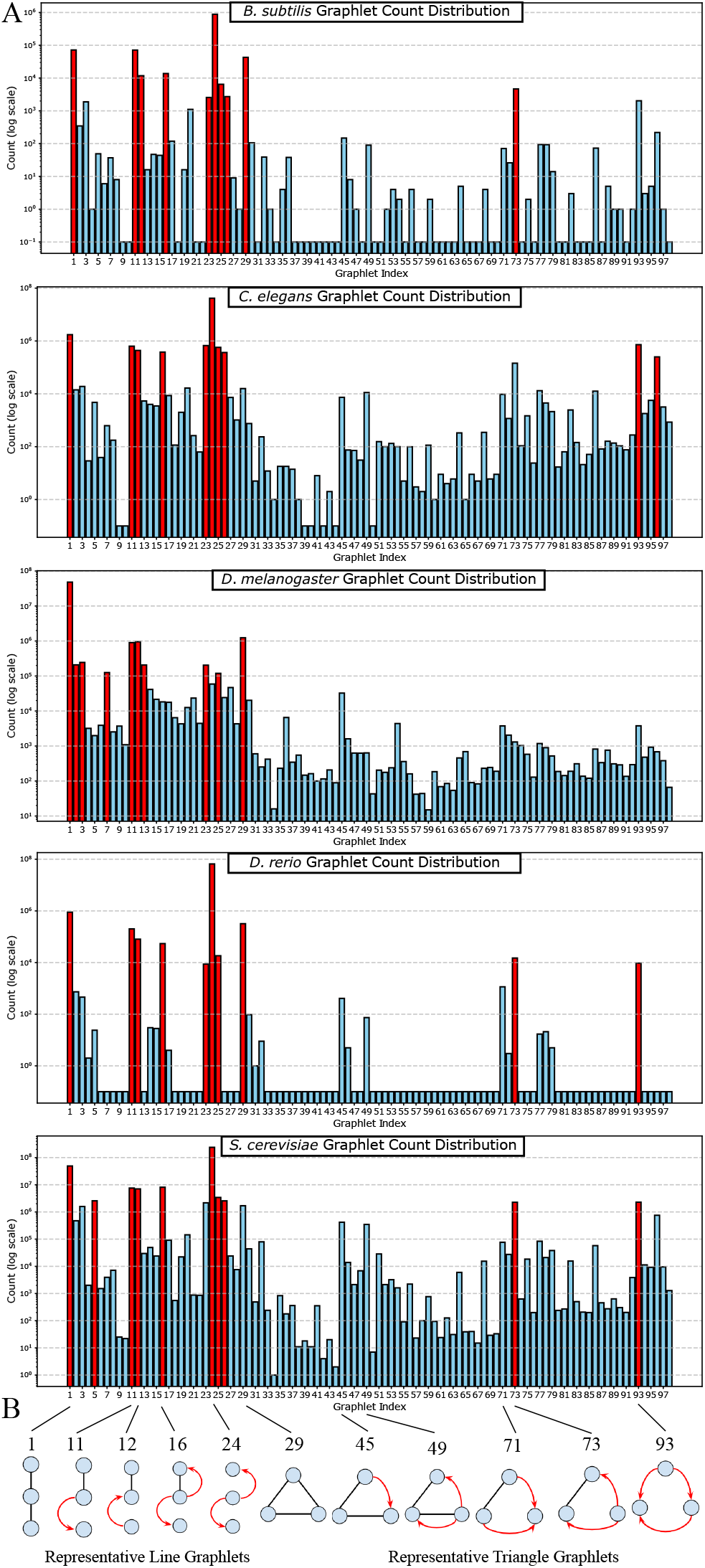
RPI Graphlet characterizations of the five species networks. (A) Graphlet count distributions of the 98 graphlets for each species network (red bars indicate the 10 most frequent graphlets). (B) Representative 3-node graphlets that appear frequently in at least one species.

### 3.2 Graphlets in Oxidative Stress Subnetworks

We generated oxidative stress subnetworks that connected 80% of the oxidative stress response proteins within each species network and identified frequently occurring graphlets compared to random networks (Methods). All 98 potential 3-node graphlets were present in at least one of the oxidative stress subnetworks, and 48 unique graphlets were significant in at least one subnetwork (Table 3). *S. cerevisiae* had the largest number and the most diverse array of significant graphlets out of all species, with 31 of the 96 unique graphlets found significant (*p <* 0.01) in the oxidative stress subnetwork compared to 1,000 randomized subnetworks. In contrast, *D. rerio* had the least diverse array of significant graphlets, with only three significant in the oxidative stress subnetwork compared to the randomized subnetworks. Although all graphlets were present in at least one subnetwork, only nine graphlets were present in the subnetworks of all five species: two graphlets with only physical interactions, four graphlets with only regulatory interactions, and three graphlets with mixed regulatory and physical interactions.

**Table 3.** Oxidative stress subnetwork sizes and their count of unique and significant 3-node graphlets (*p <* 0.01).

| Org. | Nodes | Edges |  | Graphlets |  |
| --- | --- | --- | --- | --- | --- |
|  |  | PPI | Reg. | Unique | Significant |
| *B. sub.* | 26 | 34 | 39 | 17 | 7 |
| *C. ele.* | 124 | 452 | 2,686 | 88 | 19 |
| *D. mela.* | 140 | 1,436 | 217 | 84 | 12 |
| *D. rer.* | 49 | 24 | 87 | 12 | 3 |
| *S. cer.* | 177 | 2,492 | 5,709 | 96 | 31 |

Traditional algorithms that operate on a uniform edge type cannot identify mixed graphlets, which make up over 80% of the graphlets we are counting. The significant mixed graphlets (*p <* 0.01) that occur most frequently differed between the oxidative stress response networks of all species (Figure 3, Supplementary Tables S4–S8). Within the most frequent mixed graphlets across species are three unique triangle-graphlets and five unique line-graphlets. All species but *S. cerevisiae* have a mixed graphlet in their top three most common significant graphlets; however, 26 of the 31 significant graphlets in *S. cerevisiae*’s oxidative stress subnetwork were mixed graphlets (Supplementary Table S8). *B. subtilis, D. melanogaster*, and *S. cerevisiae* have graphlets in which a single edge has a regulatory and physical interaction, making up a small percentage of edges in the total network. Thus, seeing it in one of the most frequently occurring significant graphlets is surprising. Again, we note a similarity among the most common significant mixed graphlets across species, since some differ only by a single edge.

**Figure 3.**
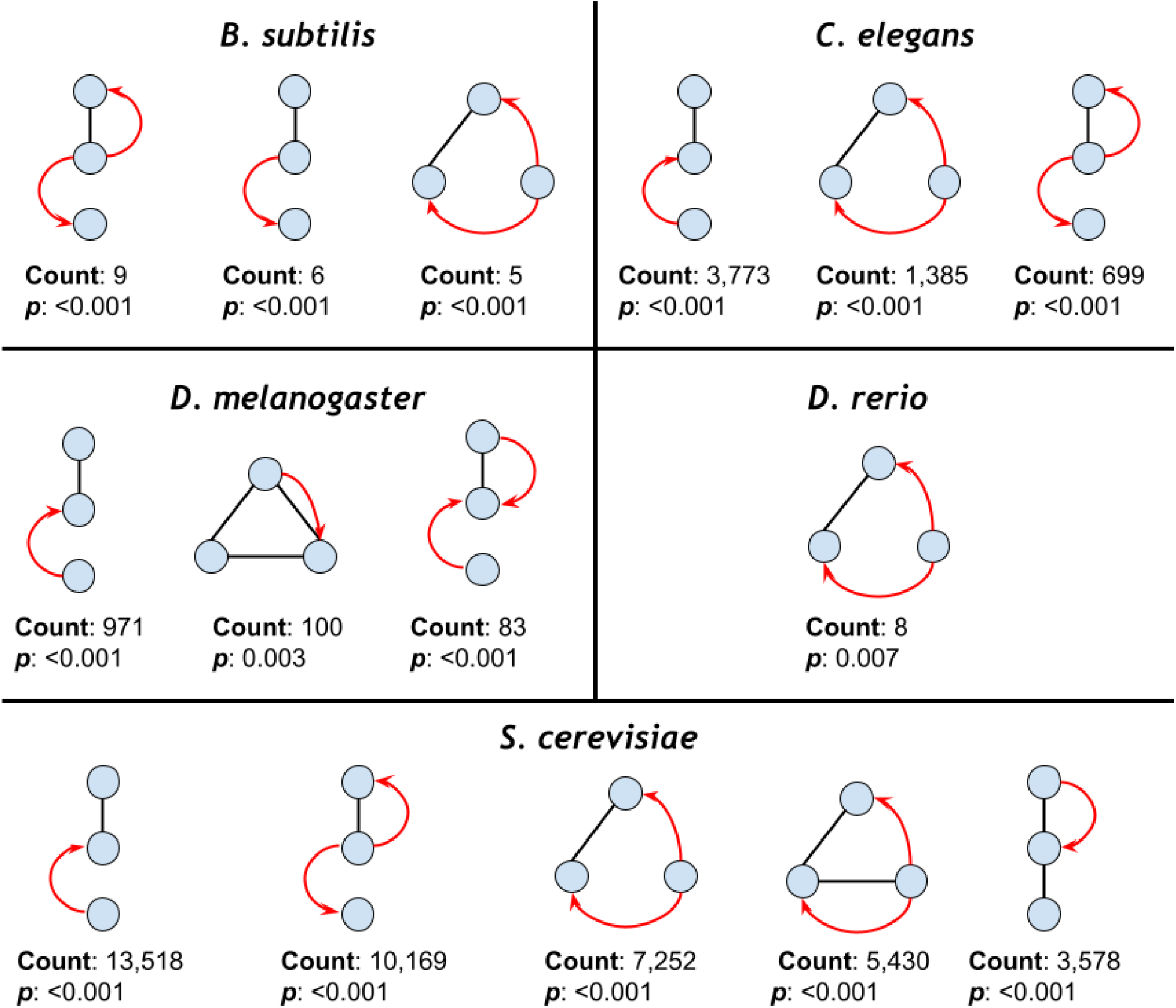
The most common mixed graphlets in each species’ oxidative stress response networks (*p <* 0.01).

None of the nine graphlets found in all subnetworks were significantly over-represented in all five species. However, five graphlets were significantly over-represented in at least three species, and two graphlets were significantly over-represented in at least four of the species. Among these five graphlets, three had mixed interactions, and two had only regulatory interactions (Figure 4). All five graphlets (including the three mixed graphlets) appeared at least once in one species’ top three most common significant graphlets.

**Figure 4.**
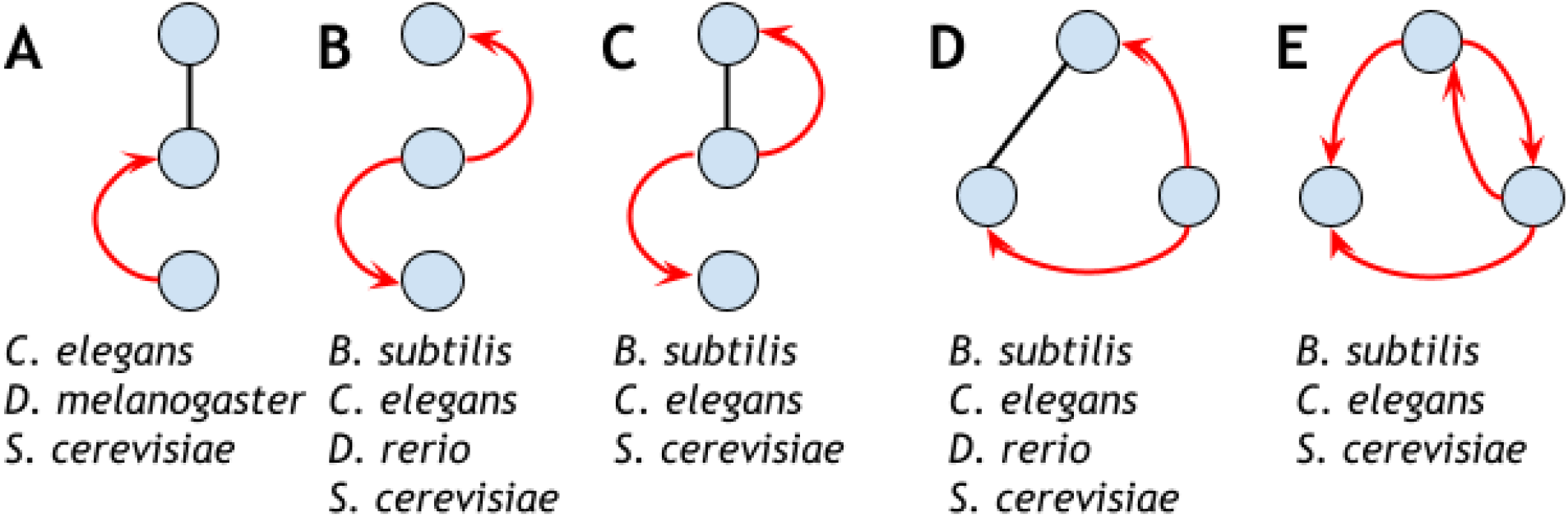
The five 3-node RPI graphlets found significant in at least three species’ oxidative stress response networks.

Interestingly, the only significant mixed graphlet in *D. rerio* also appears among the most common significant mixed graphlets in *B. subtilis, C. elegans*, and *S. cerevisiae* (Figure 3 and Figure 4D). This graphlet outlines a transcription factor that regulates the genes of two proteins that physically interact together. As a case study, we visualized this graphlet within the *B. subtilis* (Figure 5) and *D. rerio* (Supplementary Section S3) oxidative stress subnetworks. The graphlet of interest represents a transcription factor that regulates two genes whose protein products physically interact with each other, a pattern that neither physical interactions nor regulatory interactions alone would capture.

**Figure 5.**
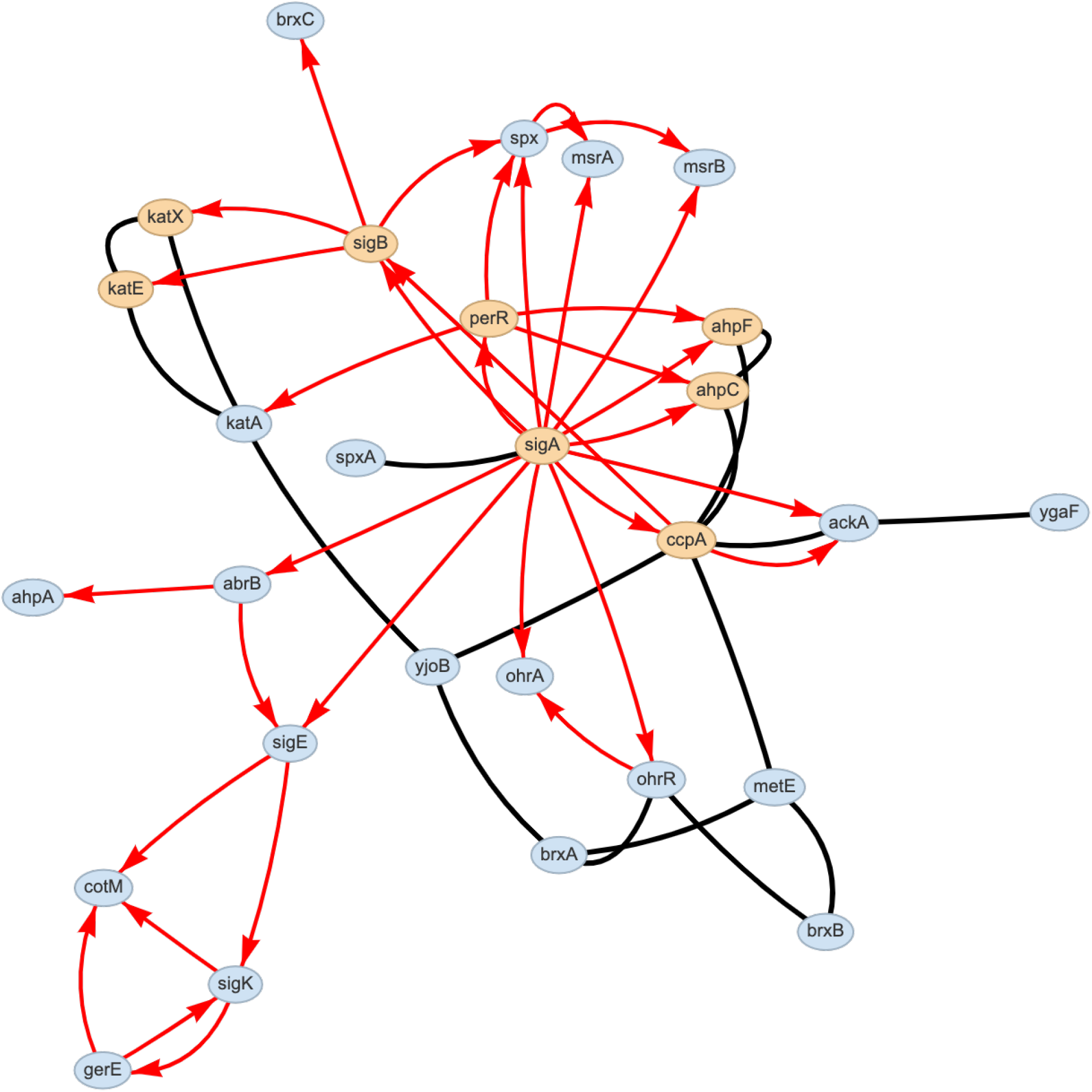
*B. subtilis* oxidative stress response network with the nodes involved in the mixed graphlet over-represented in four species highlighted in yellow (Figure 4D).

The *B. subtilis* oxidative stress subnetwork contains RNA polymerase sigma factors SigA, which initiates transcription in all growth phases, under stress, and during sporulation, and SigB, which is involved in stress response [27]. SigA regulates the genes *ahpC, ahpF*, and *ccpA*, whose protein products physically interact with each other. The AhpC and AhpF proteins form an alkyl hydroperoxide reductase (AhpCF) that detoxifies free radicals under conditions of oxidative stress [18]. The CcpA protein (catabolite control protein A) regulates genes under conditions of excess carbon and is important for survival under conditions of acidic or oxidative stress [28].

The alternative sigma factor, SigB, regulates the genes *katE* and *katX*, whose proteins physically interact. KatE, also known as KatB, is a catalase specifically regulated by the stress-dependent sigma factor SigB, while KatX is a catalase expressed only during sporulation in *B. subtilis* [6]. The last graphlet represents the AhpCF complex regulated by a hydrogen and peroxide-activated transcription factor, PerR, a known activator of the AhpCF complex [18].

### 3.3 Orbits in Oxidative Stress Proteins

We then identified specific orbits that tend to be occupied by proteins related to oxidative stress for each species. Significantly over-represented orbits may indicate that there is a particular role the oxidative stress proteins play within the network. We found 0 significant orbits in *B. subtilis*, 9 significant orbits in *C. elegans*, 2 significant orbits in *D. rerio*, 22 significant orbits in *D. melanogaster*, and 48 significant orbits in *S. cerevisiae* (*p <* 0.01 with 1,000 random samples; Methods). Since *B. subtilis* has no significant orbits (presumably because there were only 18 oxidative stress proteins), we focus on the remaining four species for the rest of this section. Two of the significant orbits, *O*_1_ and *O*_78_, were common to all four species, and four of the significant orbits are common to *C. elegans, D. melanogaster*, and *S. cerevisiae*. Three of these four significant orbits were mixed orbits, *O*_29_, *O*_30_, and *O*_31_ (Figure 6). We also found sets of significant orbits that formed complete graphlets (Figure 7).

**Figure 6.**
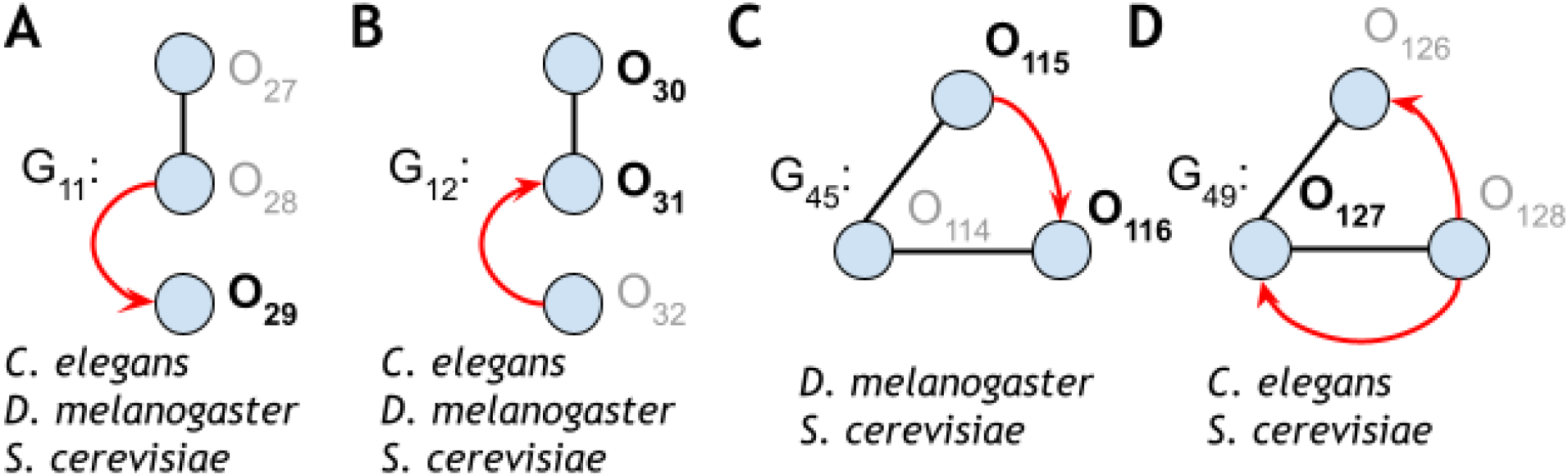
Mixed graphlets that had at least one significant orbit position in more than one species (*p <* 0.01). Orbits significant across the species listed are highlighted in bold.

**Figure 7.**
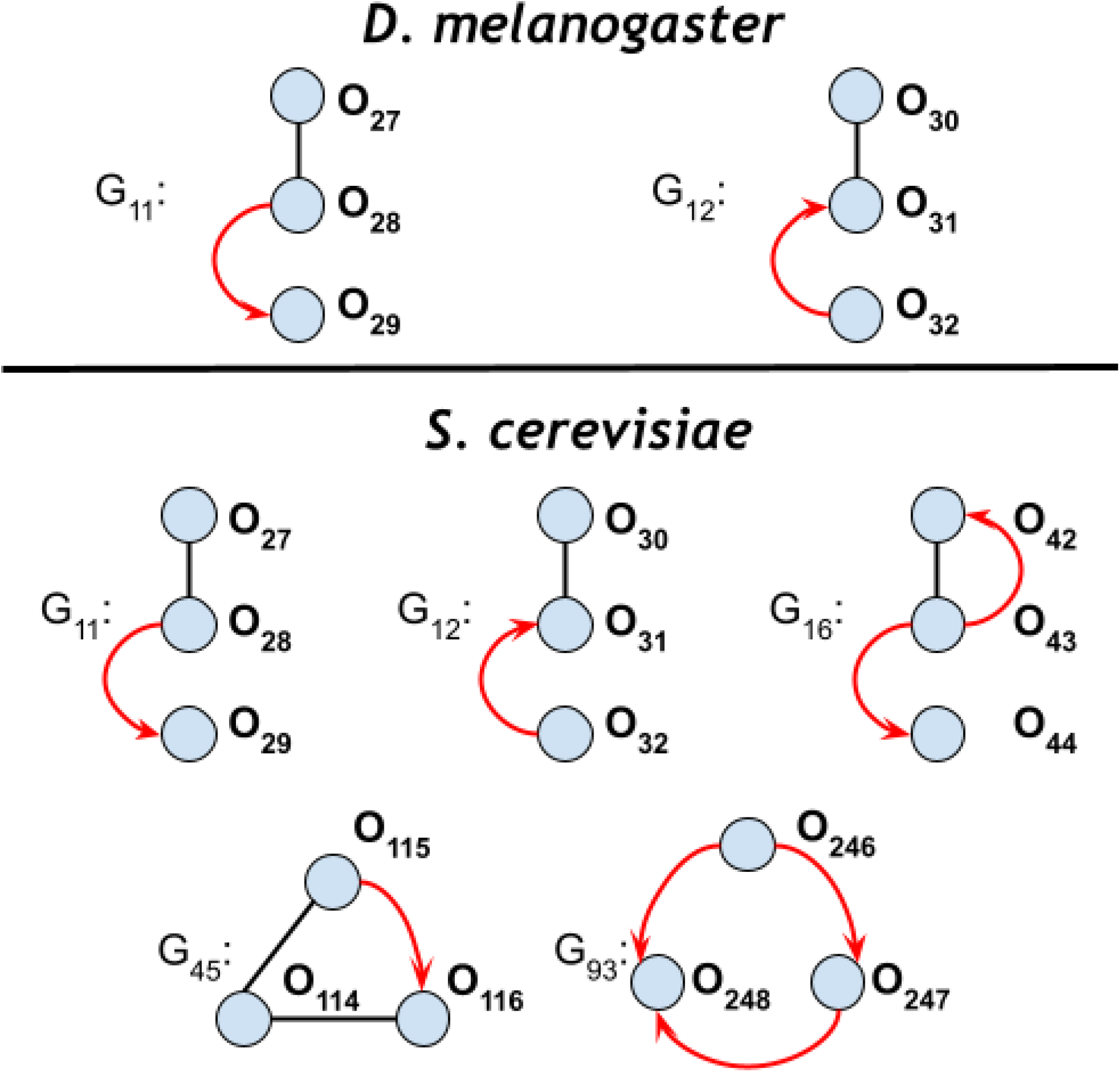
RPI graphlets where all three orbits in the graphlet were over-represented (*p <* 0.001 for all orbits shown).

To better understand the potential role of specific significant orbits, we computed the Gene Ontology enrichment of the oxidative stress proteins that participated in each significant orbit in each species (Methods). Again, we focused on significant orbits from mixed graphlets, since these had never been previously computed. Our GO enrichment analysis revealed that significant orbits in three out of the five species studied contained significantly overrepresented GO terms: *C. elegans, D. melanogaster*, and *S. cerevisiae*. Most enriched terms fell under the biological process ontology (Supplementary Figure S12).

Finally, we wanted to ask whether orbit over-enrichment provided additional information beyond graphlet over-enrichment. Here, we focus the orbit *O*_115_ in *S. cerevisiae* as an interesting, significant mixed orbit within the graphlet *G*_45_ (Figure 6C). *G*_45_ was not found to be significantly over-represented in the *S. cerevisiae* graphlet-based analysis. In the orbit-based analysis above, we identified 70 enriched GO terms for *O*_115_ (Supplementary Figure S13).

We compared the 70 enriched GO terms to GO enrichment results from an induced subnetwork of *S. cerevisiae* stress response proteins that participate in *G*_45_. These nodes may occupy orbits *O*_114_ −−*O*_116_, but they require that all nodes in the graphlet are stress response proteins. Graphlet *G*_45_ had 23 enriched GO terms, all of which were found to be enriched in *O*_115_. In total, we identified 47 GO terms in the orbit-based analysis that were not identified in the graphlet-based analysis (Supplementary Figure S13). These results highlight the ability of an orbit-specific resolution to identify new functions for proteins in specific roles compared to graphlet over-representation.

## 4 Discussion

GRPhIN is a new graphlet-counting approach for enumerating mixed patterns of undirected physical and directed regulatory interactions. GRPhIN adopts a graphlet and orbit encoding scheme that breaks isomorphic (graphlet) and automorphic (orbit) symmetries in a two-step sorting process (Figure 1). We applied GRPhIN to the networks of five species, identifying commonly abundant graphlets despite their varying size and composition of physical and regulatory interactions (Figure 2). We then identified graphlets that were significantly over-represented in oxidative stress subnetworks across all species (Table 3), highlighting the graphlets with mixed physical and regulatory interactions (Figure 3). To our knowledge, no other publicly-available tool enumerates these mixed graphlets.

We also identified graphlets that were significant in at least three species’ oxidative stress subnetworks (Figure 4). Furthermore, we illustrated a graphlet that describes a transcription factor that regulates two physically interacting proteins in *B. subtilis* (Figure 5) and *D. rerio* (Supplementary Section S3). Additionally, we focused on identifying significant orbits occupied by proteins related to the oxidative stress response and found significant orbits present in four of the five species. Then, we identified multiple examples of mixed graphlets with at least one significant orbit in more than one species (Figure 6). Finally, we illustrated how orbit-specific over-representation can identify different biological processes than a graphlet over-representation.

GRPhIN sorts encoded edge types in an enumerative algorithm, which can count graphlets and orbits quickly in practice (the largest network took less than 20 minutes on a MacBook Pro 2024, Table 1). However, there are many opportunities to optimize this algorithm, especially by adopting an analytic approach to graphlet counting similar to ORCA [14, 26]. Increasing the graphlet size to consider 4-node and 5-node graphlets will require a new approach for enumerating mixed graphlets. While all graphlets appeared at least once across the species networks, an overwhelming number of them occur with very low frequency in some species (Figure 2), presenting an opportunity for a greedy approach to count larger graphlets.

Although GRPhIN has revealed similarities among the five species networks, additional work is needed to identify graphlets common across taxa. The species networks are all in different states of completeness, largely due to the nature of research with those organisms. Further, graphlets have been criticized as being sensitive to network perturbations, though other work has reaffirmed their robustness and stability for describing protein interaction networks [12]. Accounting for the density and completeness of the species networks when identifying graphlets common between species remains future work.

Oxidative stress response is conserved across species [17], and our results highlight roles that oxidative stress proteins may play within the larger network. Ongoing work includes biologically characterizing over-represented graphlets and over-represented orbits found in the oxidative stress subnetworks, such as the graphlet case study we describe in Figure 5 and the orbit-specific over-representation analysis of *O*_115_ in graphlet *G*_45_ in *S. cerevisiae*. Identifying the roles of key proteins within mixed RPI graphlets could provide insight into the coordination behind oxidative stress responses beyond single network types. In addition, further comparison across species could identify whether specific stress-response graphlets are universal or organism-specific. Oxidative stress is only one type of stress, and other stress response pathways may be universally conserved, further analysis into different oxidative stress subnetworks could provide insights into whether graphlets are specific to oxidative stress responses or are enriched in other stress networks. Lastly, the GO enrichment analysis on a set of oxidative stress related proteins at an orbit could be further analyzed to look at both cross species and orbit specific configurations.

## 6 Author contributions statement

A.R. conceived the project and supervised all work; A.B. and O.A. implemented all code and conducted all experiments. All authors wrote and reviewed the manuscript.

## 7 Acknowledgments

This work is supported by the National Science Foundation (NSF: #1750981).

