## Supplementary Information for "GRPhIN: Graphlet Characterization of Regulatory and Physical Interaction Networks"

### *Supplementary Information for*
### GRPhIN: Graphlet Characterization of Regulatory and Physical Interaction Networks

Altaf Barelvi[1,†], Oliver Anderson[1,†], and Anna Ritz[1,*]

[1]Biology Department, Reed College, Portland, Oregon, USA
[†]These authors contributed equally

#### Contents

**S1 Species Data Sources** 2

**S2 Enumerated Graphlets** 3

**S3 Graphlet Distribution Across Species** 5

**S4 Random Walk with Restart for Oxidative Stress Networks** 7

**S5 Degree-Preserving Randomized Networks** 7

**S7 Orbit Enrichment in Oxidative Stress Response Proteins** 19

#### S1 Species Data Sources

GRPhIN contains physical and regulatory information for five different species sourced from ProteinWeaver, an interactive network visualization tool that connects proteins to biological functions of interest within a molecular interaction network [3]. For completeness, we provide the same information below.

***B. subtilis***: To generate the PPI network for bacteria *B. subtilis*, we exported the "Interaction" dataset from SubtiWiki (v4) [22] and merged it with experimental, text-mined, and database-validated interactions from STRING-DB's (v12) [28] "Physical Links" dataset. To generate the GRN, we exported the "Regulatory" dataset from SubtiWiki (v4). In addition, Gene Ontology (GO) annotations were obtained from the "Gene" data set from the SubtiWiki (v4) export page and the QuickGO annotation service from EBI [5]. UniProt IDs [2] were assigned using the AmiGO 2 search function on the GO web server [6].

***C. elegans***: To generate the PPI network for the nematode *C. elegans* we exported the WormBase (v18) [27] interactome and filtered it to only physical interactions where both proteins were verified by UniProt [2]. To generate the GRN, regulatory interactions were exported from TFLink [19]. GO term association data were also obtained through the WormBase database, and we used the UniProt namespace mapping service to standardize WormBase IDs to UniProt IDs.

***D. melanogaster***: To generate the PPI network for the fruit fly *D. melanogaster*, we exported an existing fly PPI network from a compilation of six fly interaction databases [32]. Most of the interactions in the PPI network originate from the Drosophila Interactions Database (DroID) [20]. To generate the GRN, we exported the "Genetic Interaction Table" dataset from FlyBase (v2024_03) [10, 33]. For the GO term association data, we exported FlyBase's Gene Association file and merged it with a dataset generated from the EBI's QuickGO annotation service [5]. FlyBase IDs were standardized to UniProt IDs using the UniProt name mapping service [2].

***D. rerio***: To generate the PPI network for *D. rerio*, we scraped XML data from the PSICQUIC database [8] with a Python script. With JavaScript, we pre-processed the XML into JSON, and with an online converter [1], we converted the JSON into a CSV. We also downloaded protein aliases and interactions from STRING-DB (v12) [28]. We filtered the STRING-DB data to include only experimental, text-mined, and database-validated interactions. We merged the PSICQUIC and STRING-DB data together to form the final dataset. In addition, we standardized the protein identifiers to UniProt identifiers via the UniProt ID mapping service [2]. During this process, we found that many of the protein identifiers scraped from PSICQUIC were obsolete. Thus, we filtered the PPI data to only UniProt-verified proteins. To generate the GRN, we exported regulatory interactions from TFLink [19]. To get the GO association data for *D. rerio*, we used EBI's QuickGO service [4].

***S. cerevisiae***: To generate the PPI for brewer's yeast (*S. cerevisiae*), we downloaded the PPI network from BioGRID [21]. To generate the GRN network for *S. cerevisiae*, we exported regulatory interactions from TFLink's database [19]. To get the GO term annotation data, we used QuickGO to find all reviewed annotations [4]. We used the UniProt namespace mapping service to standardize BioGRID IDs to UniProt IDs [2].

The scripts used to process and generate these datasets are available in the ProteinWeaver GitHub repository `https://github.com/Reed-CompBio/protein-weaver/` [3].

### S2 Enumerated Graphlets

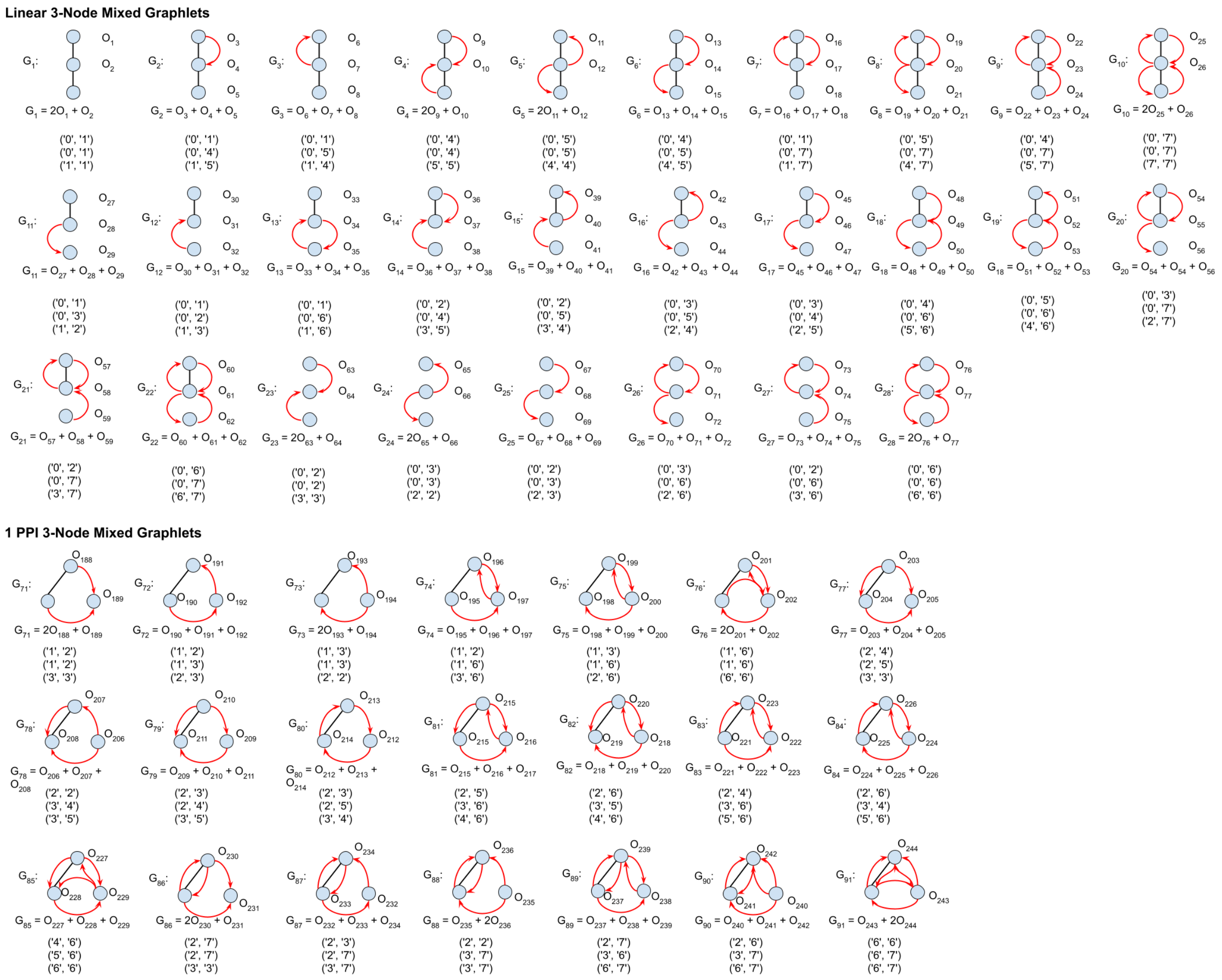

Figure S1: All RPI graphlets and graphlet encodings (along with Figure S2). Graphlets are numbered $G_1$ through $G_{98}$ and orbits are numbered $O_1$ through $O_{259}$.

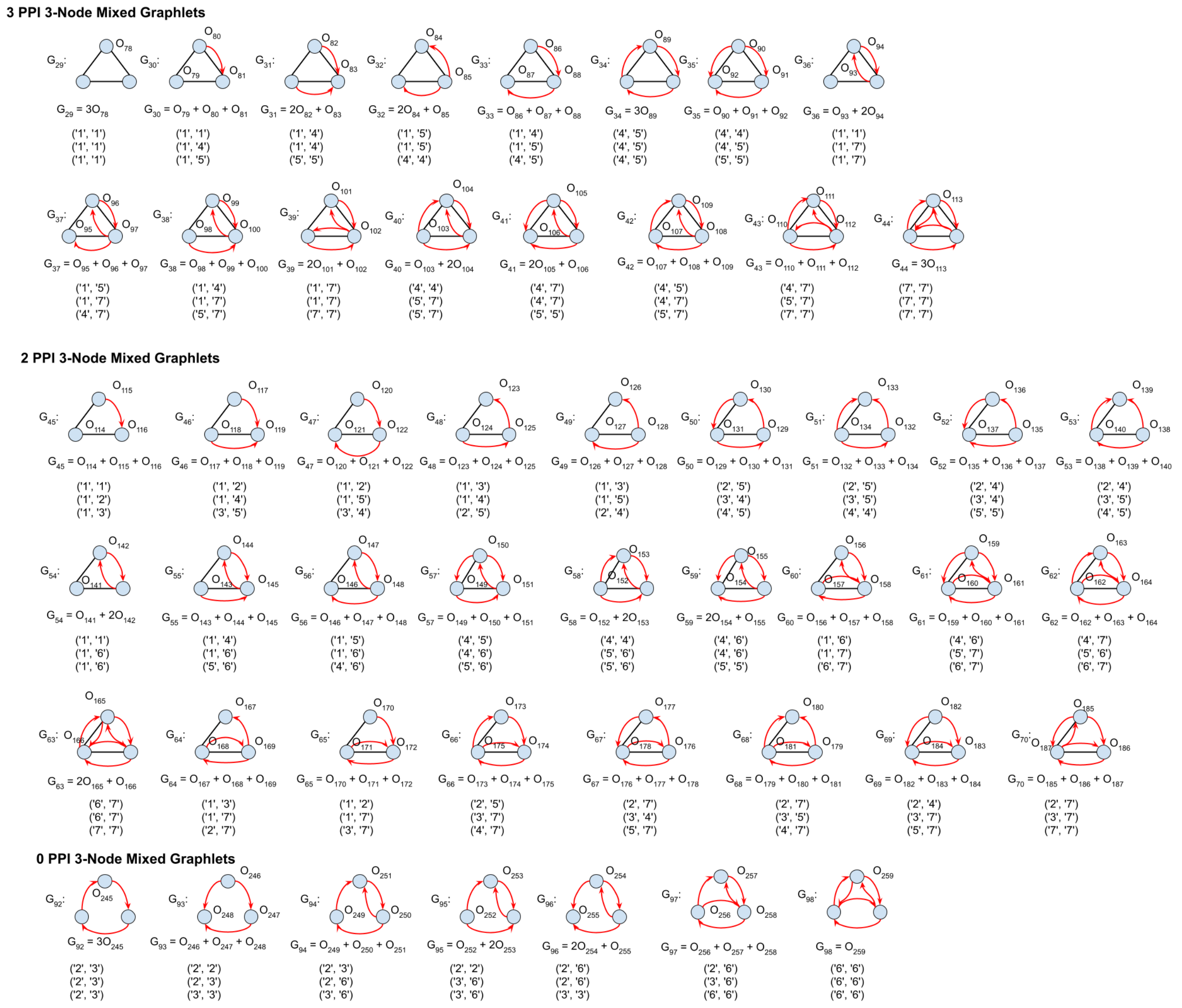

Figure S2: All RPI graphlets and graphlet encodings (along with Figure S1). Graphlets are numbered $G_1$ through $G_{98}$ and orbits are numbered $O_1$ through $O_{259}$.

### S3 Graphlet Distribution Across Species

| Species | Total Graphlets | Total Mixed % | Total Graphlets w/ Filter | Total Mixed % w/ Filter |
|---|---|---|---|---|
| *B. sub.* | 1,117,428 | 9.5 | 162,963 | 65.1 |
| *C. ele.* | 47,653,925 | 3.7 | 4,366,464 | 40.0 |
| *D. mela.* | 52,672,308 | 5.5 | 4,535,268 | 63.8 |
| *D. rer.* | 67,288,323 | 0.5 | 709680 | 49.8 |
| *S. cer.* | 331,323,465 | 9.4 | 44,250,332 | 70.8 |

Table S1: The total graphlet counts and total mixed graphlet percentage in two conditions across five species (*B. subtilis*, *C. elegans*, *D. melanogaster*, *D. rerio*, and *S. cerevisiae*). The second and third columns include all graphlets, and the fourth and fifth columns remove $G_1$ and $G_{24}$. These two graphlets are a PPI-only line graphlet and a regulatory-only line graphlet, which together account for a large proportion of graphlets in the five species networks.

| Species | Line % | Triangle % | Line % w/ Filter | Triangle % w/ Filter |
|---|---|---|---|---|
| *B. sub.* | 99.3 | 0.7 | 95.2 | 4.8 |
| *D. rer.* | 99.9 | 0.1 | 96.3 | 3.8 |
| *D. mela.* | 99.8 | 0.2 | 97.9 | 2.1 |
| *C. ele.* | 97.4 | 2.6 | 72.5 | 27.5 |
| *S. cer.* | 98.0 | 2 | 84.9 | 15.1 |

Table S2: Percentage of line graphlets and triangle graphlets across all five species (*B. subtilis*, *C. elegans*, *D. melanogaster*, *D. rerio*, and *S. cerevisiae*).

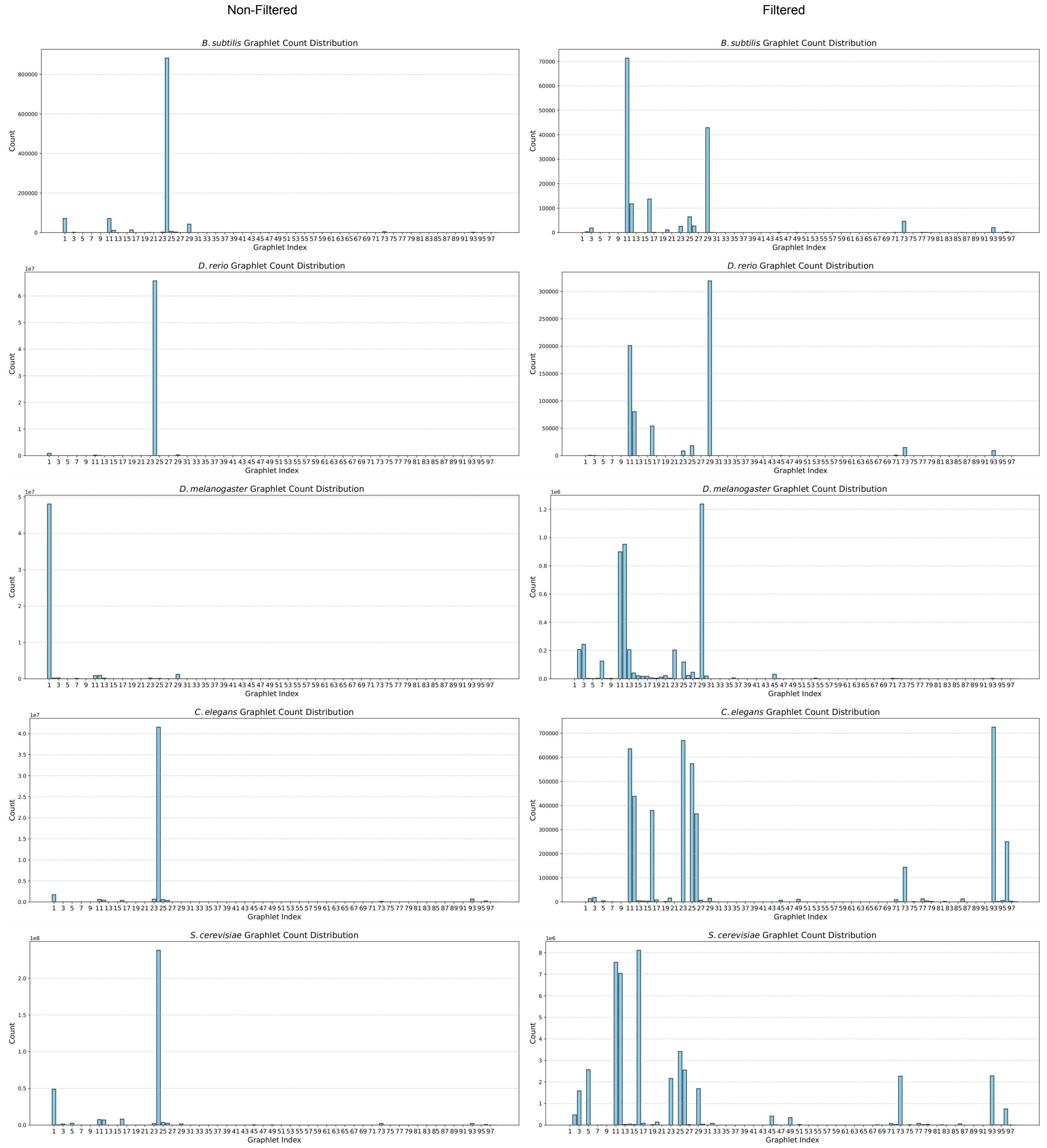

Figure S3: Graphlet distributions across the five species. Left column are the distributions where all the graphlets are included. Right column filters out $G_1$ and $G_{24}$.

#### S4 Random Walk with Restart for Oxidative Stress Networks

Random Walk with Restart (RWR) was performed to reduce the number of individual components in the induced subnetwork generated by the list of oxidative stress proteins for each species (Table S3). *D. melanogaster* and *D. rerio* have more than one connected component in their RWR networks. These consist of a large component with most of the nodes and individual nodes in isolation: likely stress proteins that the random walk restarted at but did not connect to other nodes in the subnetwork.

| | Number of Connected Components | |
|---|---|---|
| **Organism** | **Induced Subnetwork** | **RWR Subnetwork** |
| *B. subtilis* | 12 | 1 |
| *C. elegans* | 26 | 1 |
| *D. melanogaster* | 49 | 4 |
| *D. rerio* | 38 | 7 |
| *S. cerevisiae* | 1 | 1 |

Table S3: The number of connected components in the induced subnetwork from the oxidative stress proteins compared to the number of connected components in the subnetwork generated by RWR.

#### S5 Degree-Preserving Randomized Networks

##### S1 Network Randomization with Edge Swapping

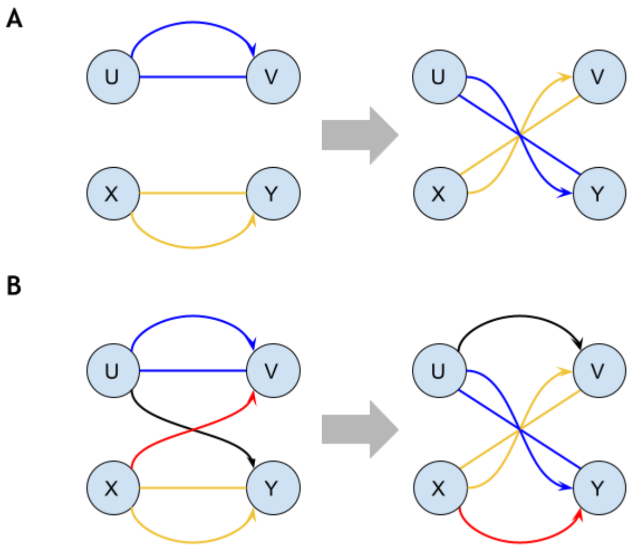

Figure S4: The switch condition for a set of four distinct nodes before an edge swap can occur during network randomization. If the "switch condition" is met, edge $uv$ (blue) can become edge $uy$ (black), and edge $xy$ (yellow) can become edge $xv$ (red) and vice versa.

For each species, we generated a random network by using the edge-swapping algorithm on the species-specific stress subnetworks. An edge swap can only occur during randomization when four unique nodes meet a "switch condition." Consider a set of four distinct nodes: $u, v, x, y$, the "switch condition" is met for these four nodes if the edges $uv$ and $xy$ are the same, and edges $uy$ and $xv$ are the same (Figure S4B) or do not exist (Figure S4A). When nodes $u, v, x, y$ meet the "switch condition," edges $uv$ and $uy$ switch, and edges $xy$ and $xv$ switch, preserving all edges and ensuring that each node's inward and outward degree stays the same after randomization.

We selected a different number of swaps for each species to ensure that there were sufficient edge swaps to properly randomize the network. We found that after a certain number of iterations, the percent difference from the original subnetwork did not increase with the number of iterations (Figures S5 and S6). We used the following iteration thresholds: 500 iterations for *B. subtilis*, 10,000 iterations for *C. elegans*, 5,000 iterations for *D. melanogaster*, 500 iterations for *D. rerio*, and 25,000 iterations for *S. cerevisiae*. *B. subtilis* and *D. rerio* had comparable randomization with increasing iterations. Thus, we chose the lowest threshold to speed up the analysis.

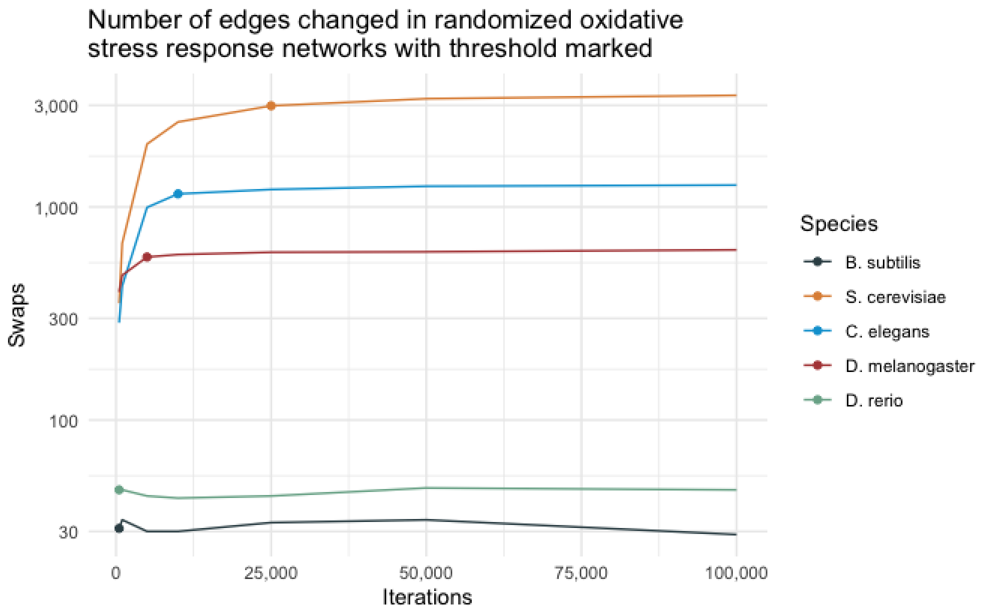

Figure S5: The number of edges changed in the randomized network compared to the original network as the number of edge-swap iterations increase.

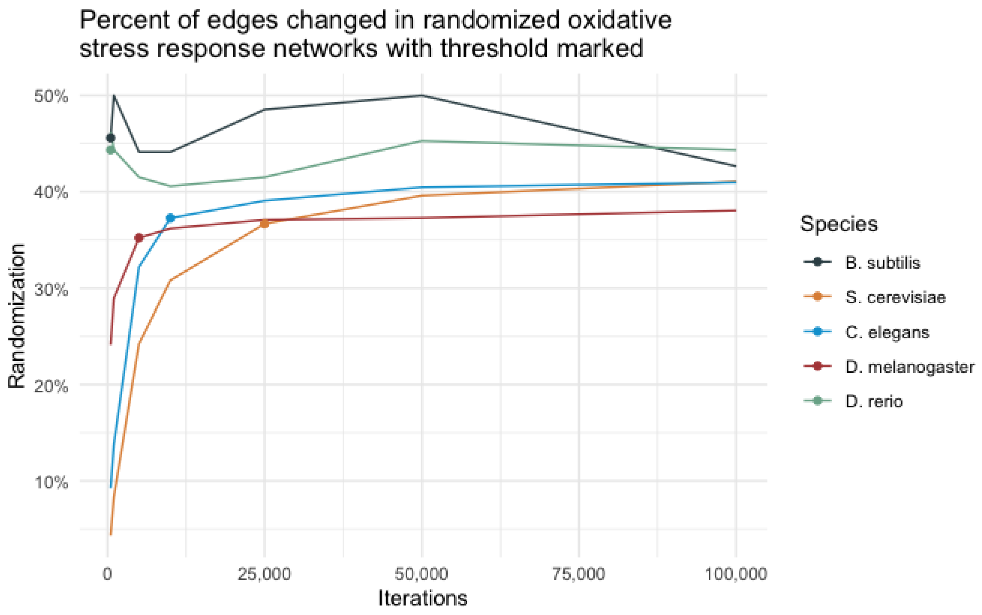

Figure S6: The percent of edges changed in the randomized network compared to the original network as the number of edge-swap iterations increase.

#### S2 Significantly Over-Represented Graphlets in Oxidative Stress Subnetworks

Each species had different graphlets that were found to be significantly over-represented in their oxidative stress subnetworks. *S. cerevisiae* had more unique significant graphlets than the others with 31 (Table S8), while *D. rerio* had the least with 3 unique significant graphlets (Table S7). *B. subtilis* had 7 graphlets that were found to be significant compared to 1,000 randomized networks (Table S4). *C. elegans* had 19 significant graphlets in its oxidative stress subnetwork (Table S5), while *D. melanogaster* had 12 significant graphlets (Table S6).

The most frequently occurring significant graphlets ($p$-value of $< 0.01$) differed between the oxidative stress response networks of all species (Figure S7). Within the top three most frequent graphlets for each species, there are ten unique graphlets. Only three unique graphlets are triangle-shaped, with one being a mixed RPI graphlet in *D. rerio*. All species but *S. cerevisiae* have an RPI graphlet in their top three most common significant graphlets.

Out of the 31 unique significant graphlets in the *S. cerevisiae* oxidative stress subnetwork, 27 were RPI graphlets. The oxidative stress subnetwork of *C. elegans* had 17 significantly over-represented RPI graphlets out of its 22 unique graphlets. In *D. melanogaster*, 12 of the 15 significantly over-represented graphlets were RPI graphlets, while in the *B. subtilis* and *D. rerio* subnetworks, there were only six and one unique RPI graphlets, respectively.

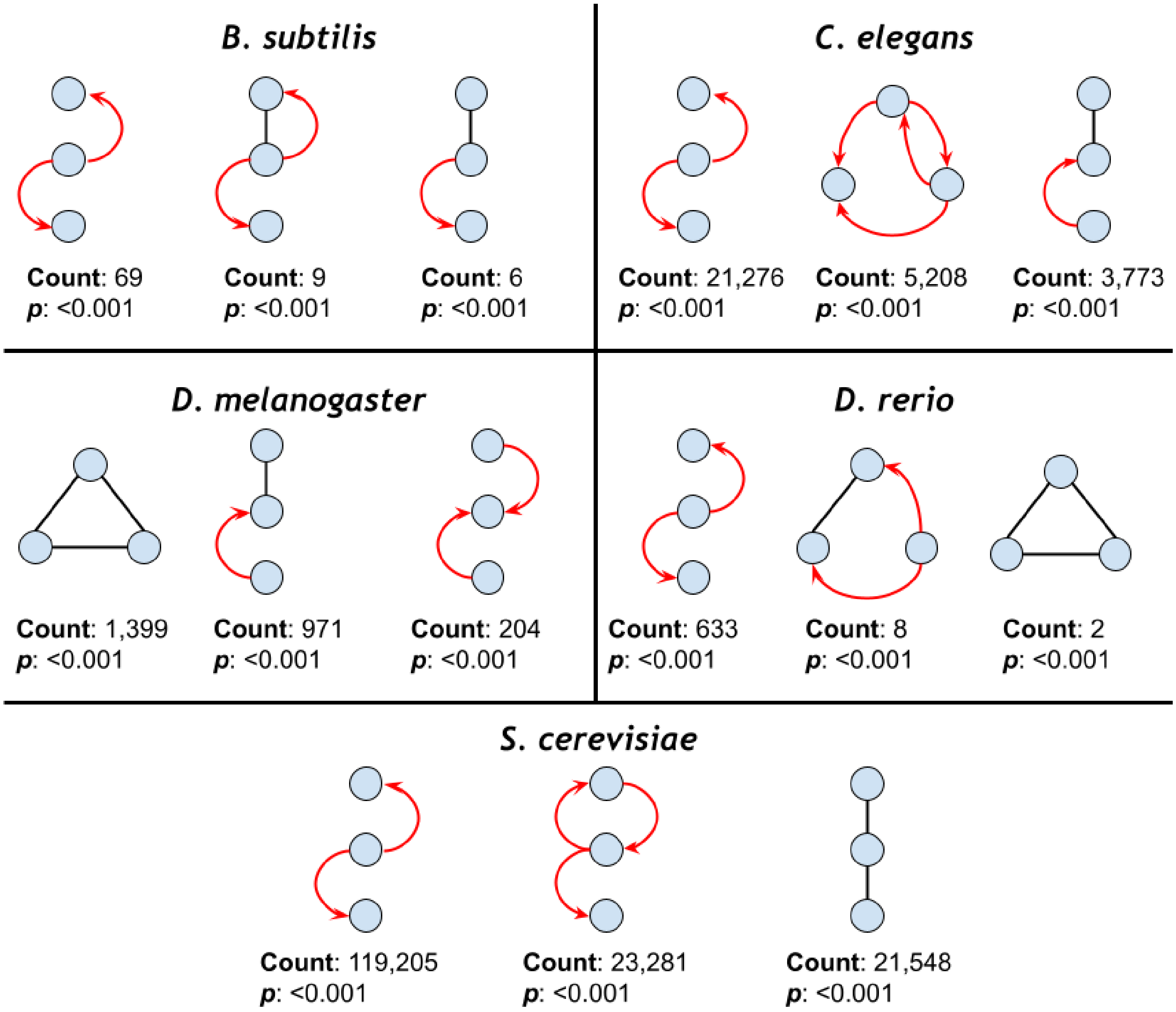

Figure S7: The three most common significant graphlets in each species' oxidative stress response networks ($p < 0.01$).

| Graphlet | Count | $p$-value |
|---|---|---|
| $G_{24}$ | 69 | $< 0.001$ |
| $G_{16}$ | 9 | $< 0.001$ |
| $G_{11}$ | 6 | $< 0.001$ |
| $G_{73}$ | 5 | 0.003 |
| $G_{3}$ | 4 | $< 0.001$ |
| $G_{77}$ | 2 | 0.006 |
| $G_{96}$ | 1 | $< 0.001$ |

Table S4: Significant graphlets in the oxidative stress response network of *B. subtilis* ($p < 0.01$). A $p$-value of $< 0.001$ indicates that the graphlet did not appear more frequently in a single randomized network than in the oxidative stress subnetwork.

| **Graphlet** | **Count** | ***p*-value** |
|---|---|---|
| $G_{24}$ | 21276 | $< 0.001$ |
| $G_{96}$ | 5208 | $< 0.001$ |
| $G_{12}$ | 3773 | $< 0.001$ |
| $G_{95}$ | 2688 | $< 0.001$ |
| $G_{73}$ | 1385 | $< 0.001$ |
| $G_{1}$ | 1378 | $< 0.001$ |
| $G_{16}$ | 699 | $< 0.001$ |
| $G_{98}$ | 693 | $< 0.001$ |
| $G_{20}$ | 344 | $< 0.001$ |
| $G_{86}$ | 312 | $< 0.001$ |
| $G_{2}$ | 255 | $< 0.001$ |
| $G_{14}$ | 255 | $< 0.001$ |
| $G_{82}$ | 224 | $< 0.001$ |
| $G_{49}$ | 181 | $< 0.001$ |
| $G_{19}$ | 163 | 0.003 |
| $G_{83}$ | 82 | $< 0.001$ |
| $G_{59}$ | 35 | $< 0.001$ |
| $G_{68}$ | 32 | $< 0.001$ |
| $G_{5}$ | 20 | 0.004 |

Table S5: Significant graphlets in the oxidative stress response network of *C. elegans* ($p < 0.01$). A $p$-value of $< 0.001$ indicates that the graphlet did not appear more frequently in a single randomized network than in the oxidative stress subnetwork.

| **Graphlet** | **Count** | ***p*-value** |
|---|---|---|
| $G_{29}$ | 1399 | $< 0.001$ |
| $G_{12}$ | 971 | $< 0.001$ |
| $G_{23}$ | 204 | $< 0.001$ |
| $G_{27}$ | 117 | $< 0.001$ |
| $G_{30}$ | 100 | 0.003 |
| $G_{14}$ | 83 | $< 0.001$ |
| $G_{21}$ | 76 | $< 0.001$ |
| $G_{15}$ | 31 | $< 0.001$ |
| $G_{88}$ | 19 | $< 0.001$ |
| $G_{18}$ | 13 | $< 0.001$ |
| $G_{38}$ | 12 | $< 0.001$ |
| $G_{46}$ | 9 | 0.005 |

Table S6: Significant graphlets in the oxidative stress response network of *D. melanogaster* ($p < 0.01$). A $p$-value of $< 0.001$ indicates that the graphlet did not appear more frequently in a single randomized network than in the oxidative stress subnetwork.

| Graphlet | Count | $p$-value |
|---|---|---|
| $G_{24}$ | 633 | $< 0.001$ |
| $G_{73}$ | 8 | 0.007 |
| $G_{29}$ | 2 | 0.005 |

Table S7: Significant graphlets in the oxidative stress response network of *D. rerio* ($p < 0.01$). A $p$-value of $< 0.001$ indicates that the graphlet did not appear more frequently in a single randomized network than in the oxidative stress subnetwork.

| Graphlet | Count | $p$-value |
|---|---|---|
| $G_{24}$ | 119205 | $< 0.001$ |
| $G_{26}$ | 23281 | $< 0.001$ |
| $G_{1}$ | 21548 | $< 0.001$ |
| $G_{96}$ | 16443 | $< 0.001$ |
| $G_{12}$ | 13518 | $< 0.001$ |
| $G_{16}$ | 10169 | $< 0.001$ |
| $G_{73}$ | 7252 | $< 0.001$ |
| $G_{49}$ | 5430 | $< 0.001$ |
| $G_{2}$ | 3578 | $< 0.001$ |
| $G_{5}$ | 1997 | $< 0.001$ |
| $G_{20}$ | 1568 | $< 0.001$ |
| $G_{28}$ | 1393 | $< 0.001$ |
| $G_{79}$ | 1370 | $< 0.001$ |
| $G_{86}$ | 1222 | $< 0.001$ |
| $G_{82}$ | 1102 | $< 0.001$ |
| $G_{19}$ | 854 | $< 0.001$ |
| $G_{75}$ | 742 | $< 0.001$ |
| $G_{32}$ | 623 | $< 0.001$ |
| $G_{51}$ | 603 | $< 0.001$ |
| $G_{56}$ | 597 | $< 0.001$ |
| $G_{68}$ | 490 | $< 0.001$ |
| $G_{52}$ | 482 | $< 0.001$ |
| $G_{59}$ | 315 | $< 0.001$ |
| $G_{54}$ | 286 | 0.007 |
| $G_{64}$ | 239 | $< 0.001$ |
| $G_{22}$ | 203 | $< 0.001$ |
| $G_{91}$ | 142 | 0.007 |
| $G_{41}$ | 68 | $< 0.001$ |
| $G_{76}$ | 57 | $< 0.001$ |
| $G_{60}$ | 43 | 0.003 |
| $G_{39}$ | 8 | 0.006 |

Table S8: Significant graphlets in the oxidative stress response network of *S. cerevisiae* ($p < 0.01$). A $p$-value of $< 0.001$ indicates that the graphlet did not appear more frequently in a single randomized network than in the oxidative stress subnetwork.

#### S3 Over-Represented *D. rerio* Graphlets in the Oxidative Stress Subnetwork

| TF | Target A | Target B |
|---|---|---|
| *perR* | *ahpC* | *ahpF* |
| *sigA* | *ahpC* | *ahpF* |
| *sigA* | *ahpC* | *ccpA* |
| *sigA* | *ahpF* | *ccpA* |
| *sigB* | *katE* | *katX* |

Table S9: Gene names of the nodes involved in the significant graphlets representing a transcription factor (TF) that regulates the genes of two proteins that physically bind one another in the oxidative stress response network of *B. subtilis*.

In addition to the *B. subtilis* oxidative stress subnetwork (Table S9), we also examined the RPI graphlet of the same TF that regulates the genes of two proteins that physically interact within the *D. rerio* oxidative stress subnetwork (Figure S8 & Table S10). We identified four unique proteins acting as TFs in the graphlet in the *D. rerio* subnetwork: FoxH1 (Fast1), GATA1a, Nanog, and Sall4. Nanog, typically associated with stem cell differentiation [26], was the TF in four of the eight unique graphlets. In one graphlet, Nanog regulates the genes for *sod1*, a Cu/Zn-superoxide dismutase, and *sod2*, a Mn-superoxide dismutase, whose protein products are known to be involved in oxidative stress response [16, 23]. In another, Nanog regulates the genes *erp44*, which encodes the endoplasmic reticulum protein 44, a thioredoxin-containing protein that is involved in the ROS-induced oxidative stress response [30], and *prdx1*, which encodes peroxiredoxin1, a thioredoxin-dependent peroxidase involved in the response to oxidative stress through the reduction of hydroperoxides [12]. In a third graphlet, Nanog regulates *keap1a* and *keap1b*, which encode Keap1a and Keap1b. Keap1 proteins are key sensors of oxidative stress that regulate the response to oxidative stress by targeting the transcription factor Nrf2 for degradation via ubiquination [18]. In the fourth graphlet, Nanog regulates the genes *keap1b* and *smad2*, a key TF in the TGF-$\beta$ pathway involved in growth and immune response. Keap1 has been shown to regulate Smad2 through physical binding [7].

| TF | Target A | Target B |
|---|---|---|
| *gata1a* | *keap1b* | *smad2* |
| *foxh1* | *keap1a* | *keap1b* |
| *nanog* | *erp44* | *prdx1* |
| *nanog* | *keap1a* | *keap1b* |
| *nanog* | *keap1a* | *smad2* |
| *nanog* | *sod1* | *sod2* |
| *sall4* | *keap1a* | *keap1b* |
| *sall4* | *keap1b* | *smad2* |

Table S10: Gene names of the nodes involved in the significant graphlets representing a transcription factor (TF) that regulates the genes of two proteins that physically bind one another in the oxidative stress response network of *D. rerio*.

Sall4, a zinc finger transcription factor, regulates the previously mentioned genes *keap1a*, *keap1b*, and *smad2*, which form two of the eight graphlets. Sall4 is involved in hematopoiesis, pectoral fin development, and maintenance of embryonic stem cells in zebrafish [11, 14]. In another graphlet,

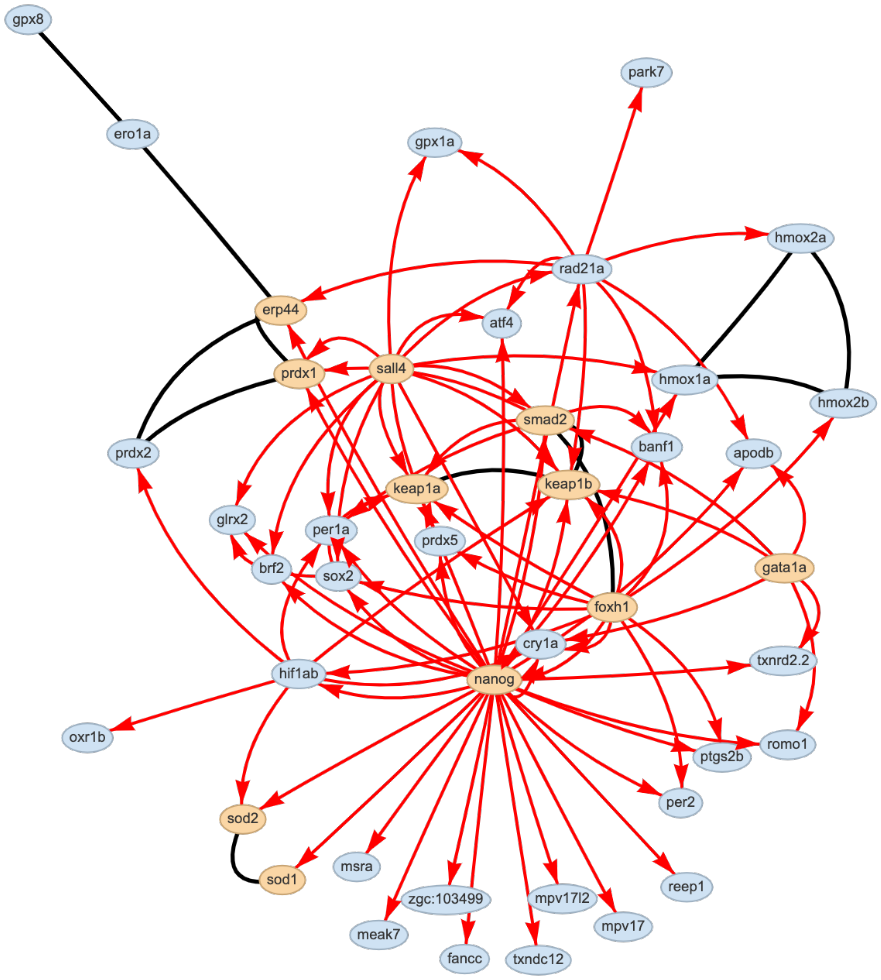

Figure S8: *D. rerio* oxidative stress response network with the nodes involved in the mixed graphlet over-represented in four species highlighted in yellow (Figure 4D).

FoxH1 regulates the genes *keap1a* and *keap1b*. FoxH1 is a transcription factor involved in the Nodal signaling pathway and structural organization during the development of zebrafish [24]. In the last graphlet, GATA1a (GATA binding protein 1a) regulates the genes encoding the proteins involved in the previously mentioned interaction between Keap1 and Smad2. GATA1a is a TF that is important during development and is involved in determining whether a cell will become an erythroid or a myeloid [9].

| Org. | Nodes | Edges PPI | Edges Reg. | Graphlets Unique | Graphlets Enriched |
|---|---|---|---|---|---|
| *B. sub.* | 26 | 34 | 39 | 17 | 3 |
| *C. ele.* | 124 | 452 | 2686 | 88 | 56 |
| *D. mela.* | 140 | 1436 | 217 | 84 | 18 |
| *D. rer.* | 49 | 24 | 87 | 12 | 4 |
| *S. cer.* | 177 | 2492 | 5709 | 96 | 72 |

Table S11: Oxidative stress response subnetwork sizes and number of unique and enriched 3-node graphlets ($\mathbb{E} > 1$ and $p < 0.05$).

### S6 Graphlet Enrichment in Oxidative Stress Response Networks

In addition to the network perturbation analysis to assess oxidative stress subnetworks, we also performed an analytical calculation to determine the probability that the number of a particular graphlet was enriched in the stress response network compared to subnetworks of the full species-specific interactome.

We note that there are limitations to this approach – namely, that the edge types in our networks do not exhibit a Poisson degree distribution, this measure may favor sparse subnetworks, and that we found through permutation tests that degree-preserving random subnetworks are rather constrained (Figure S6). While there are ways to derive equations for calculating the average number of graphlets in graphs with an arbitrary degree sequence [13], this is beyond the scope of the current work.

#### S1 Calculating Graphlet Enrichment Ratios.

Enrichment ratio scores, $\mathbb{E}$, indicate whether the number of graphlets observed in the subnetwork is surprising compared to the full interactome of a particular organism [17]. The score is the ratio of the graphlets in the subnetwork to the expected number of graphlets based on the structure of the RPI network. Enrichment ratio scores greater than one ($\mathbb{E} > 1$) suggest graphlet enrichment, while scores less than one indicate the graphlet occurs less frequently than expected. Given a graph $G = (V, E)$, a subgraph $G' = (V', E') \subseteq G$, and a graphlet $g$, let $C(G, g)$ be the count of graphlet $g$ in graph $G$. Let $|E|$ and $|E'|$ be the number of edges in the global network and subnetwork, respectively.

$$\mathbb{E} = \frac{C(G', g)}{\mu}, \text{ where} \tag{1}$$

$$\mu = \frac{|E'| C(G, g)}{|E|}. \tag{2}$$

Here, $\mu$ represents the expected number of graphlets in the subnetwork, calculated by normalizing the number of edges in the subnetwork to the global network. For all enrichment scores, we calculate $Z$-scores and use Python's `scipy` module to calculate an associated $p$-value [15, 29, 31].

#### S2 Graphlet Enrichment Results

We generated subnetworks that connected 80% of the oxidative stress response proteins within each species network (see Methods). All 98 unique 3-node graphlets were present in at least one of the oxidative stress subnetworks, and 80 unique graphlets were found to be significantly enriched in at least one subnetwork (Table S11). *S. cerevisiae* had the largest number of enriched graphlets out of all species, and it also had the most diverse array of enriched graphlets, with 72 of the 98 unique graphlets were enriched in the stress response subnetworks compared to the full RPI network. Conversely, *B. subtilis* had the lowest count of enriched graphlets in the stress response subnetwork, presumably because it was the smallest subnetwork in terms of nodes and edges. While all graphlets were present in at least one subnetwork, only nine graphlets were present in the subnetworks of all five species: two graphlets with only physical interactions, four graphlets with only regulatory interactions, and three graphlets with mixed regulatory and physical interactions.

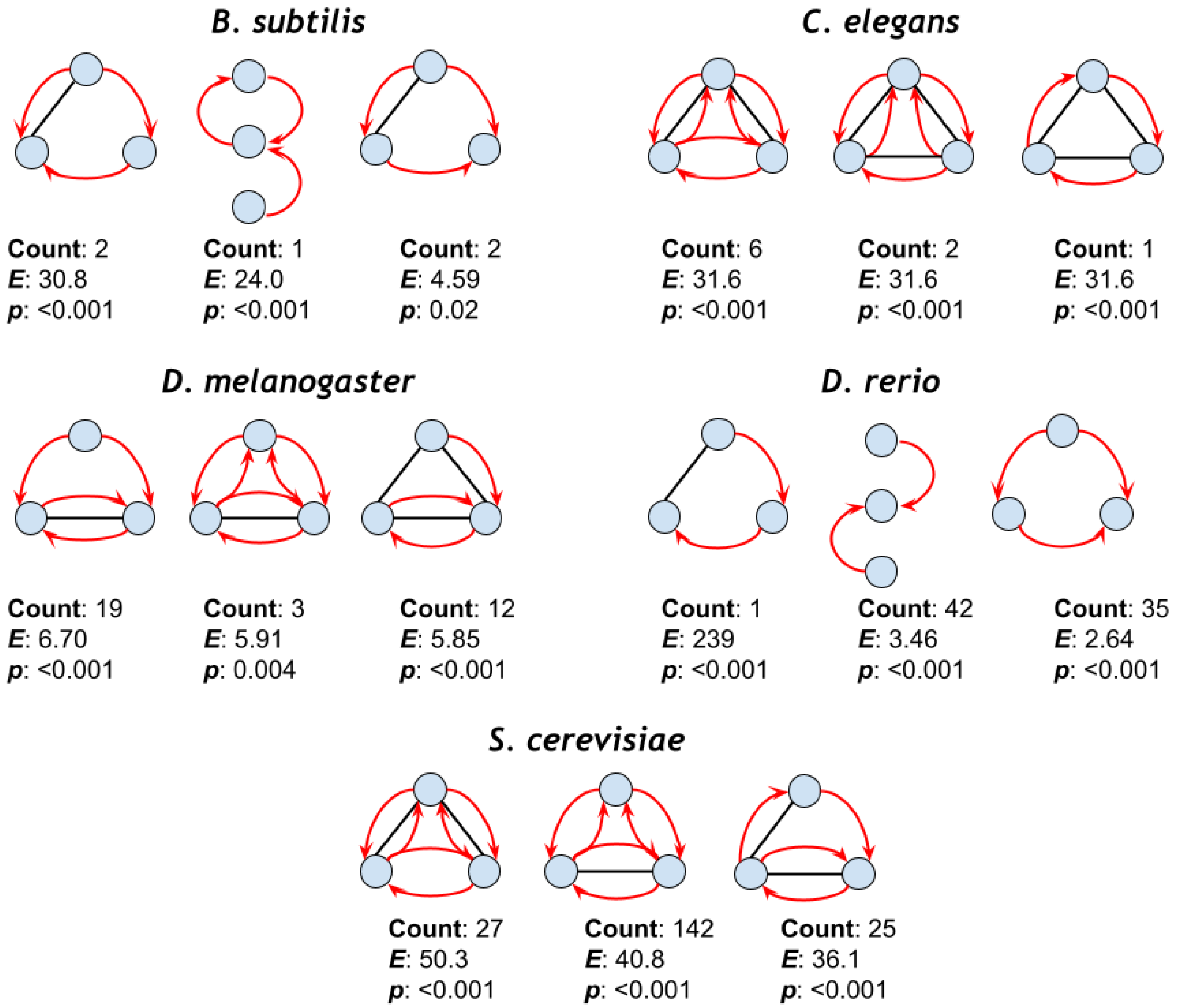

Figure S9: The three most enriched graphlets in each species' oxidative stress response networks (all have $p$-values $< 0.05$).

The graphlets most enriched ($p$ value of $< 0.05$) differed between the oxidative stress response networks of all species (Figure S9). Thirteen of the 15 enriched 3-node graphlets in Figure S9 are triangle graphlets, and 11 of the graphlets contain mixed edge types (which comprise a very small percentage of the total network, as we reported earlier). Thus, even though there are relatively few numbers of these mixed-edge-type graphlets, they are surprisingly enriched within the stress

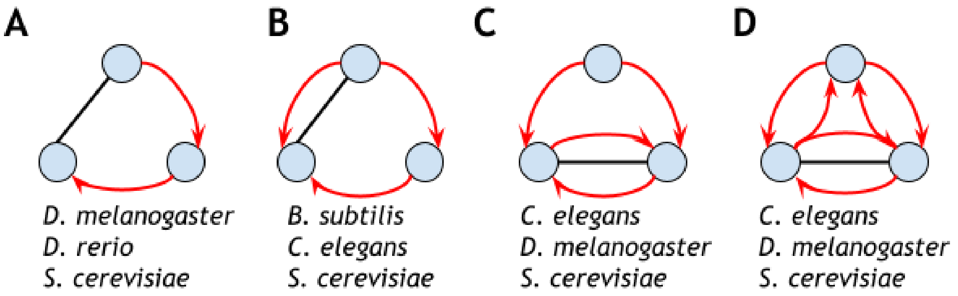

Figure S10: RPI graphlets common to at least three species' oxidative stress response networks that appear at least once in the three most enriched graphlets for each species (Figure S9).

subnetworks. We again note a similarity among the enriched graphlets, since some graphlets differ by a single edge type.

None of the nine graphlets found in all subnetworks were significantly enriched in all five species. However, 13 graphlets were significantly enriched in at least three species (including 11 graphlets with mixed interactions). Four of these mixed graphlets also appeared at least once in the top three most enriched graphlets by species (Figure S10). As a case study, we visualized the graphlet in Figure S10B within the *B. subtilis* oxidative stress subnetwork (Figure S11). The graphlet in Figure S10B is a feed-forward loop with an additional physical interaction. The *B. subtilis* oxidative stress subnetwork contains RNA polymerase sigma factor *sigA*, which initiates transcription in all growth phases, stress, and sporulation. In two mixed graphlets in *B. subtilis*, *sigA* regulates and forms a physical co-complex with *spxA*, an enzyme known to regulate growth during periods of stress [25]. In one, *sigA* regulates *perR*, a hydrogen-and peroxide-activated transcription factor that coregulates *spxA* with *sigA*. In the other, *sigA* coregulates *spxA* with *sigB*, a sigma factor that *sigA* also regulates. In these two cases, the graphlet describes a transcriptional activation complex within a feed-forward loop, a pattern that neither physical interactions nor regulatory interactions would capture alone.

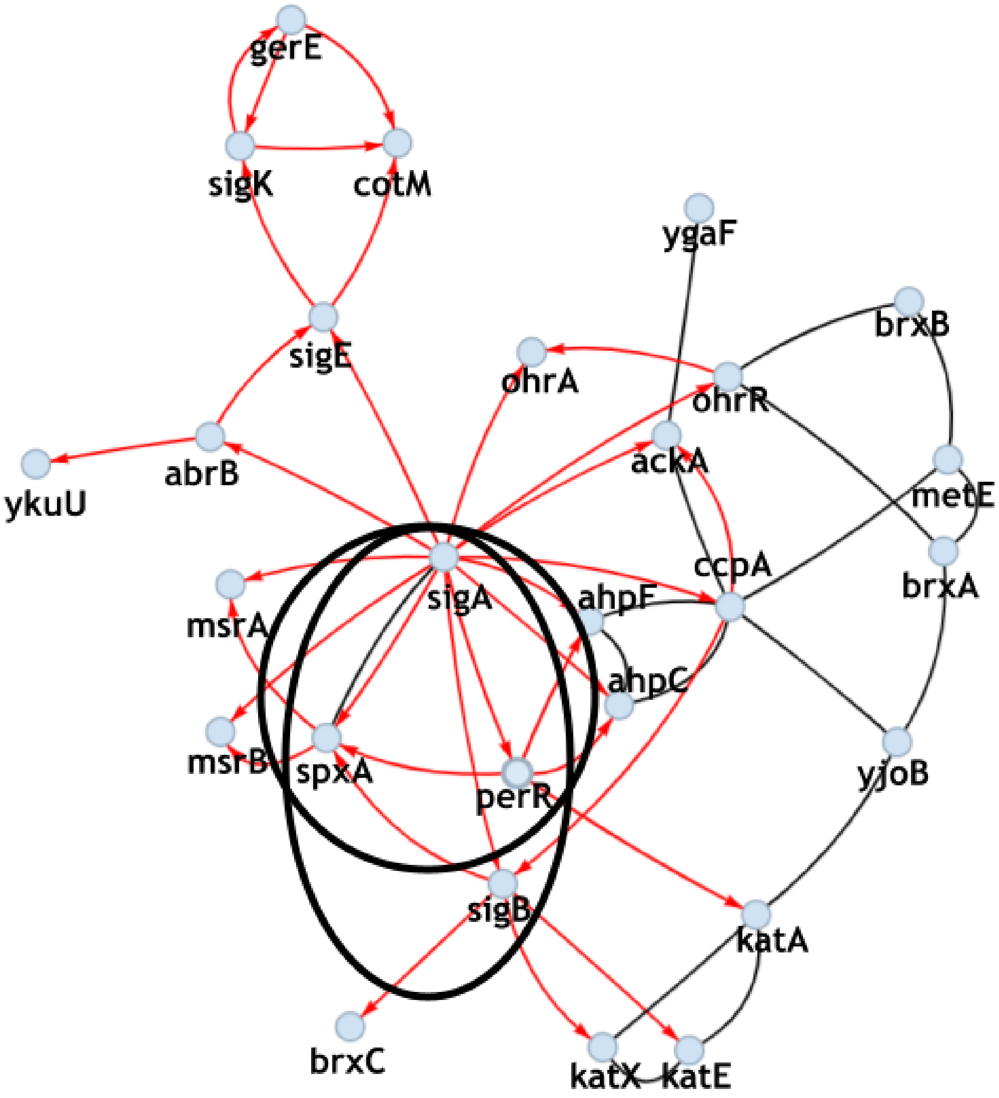

Figure S11: *B. subtilis* oxidative stress response network with the mixed graphlet enriched in three species highlighted (Figure S10B).

### S7 Orbit Enrichment in Oxidative Stress Response Proteins

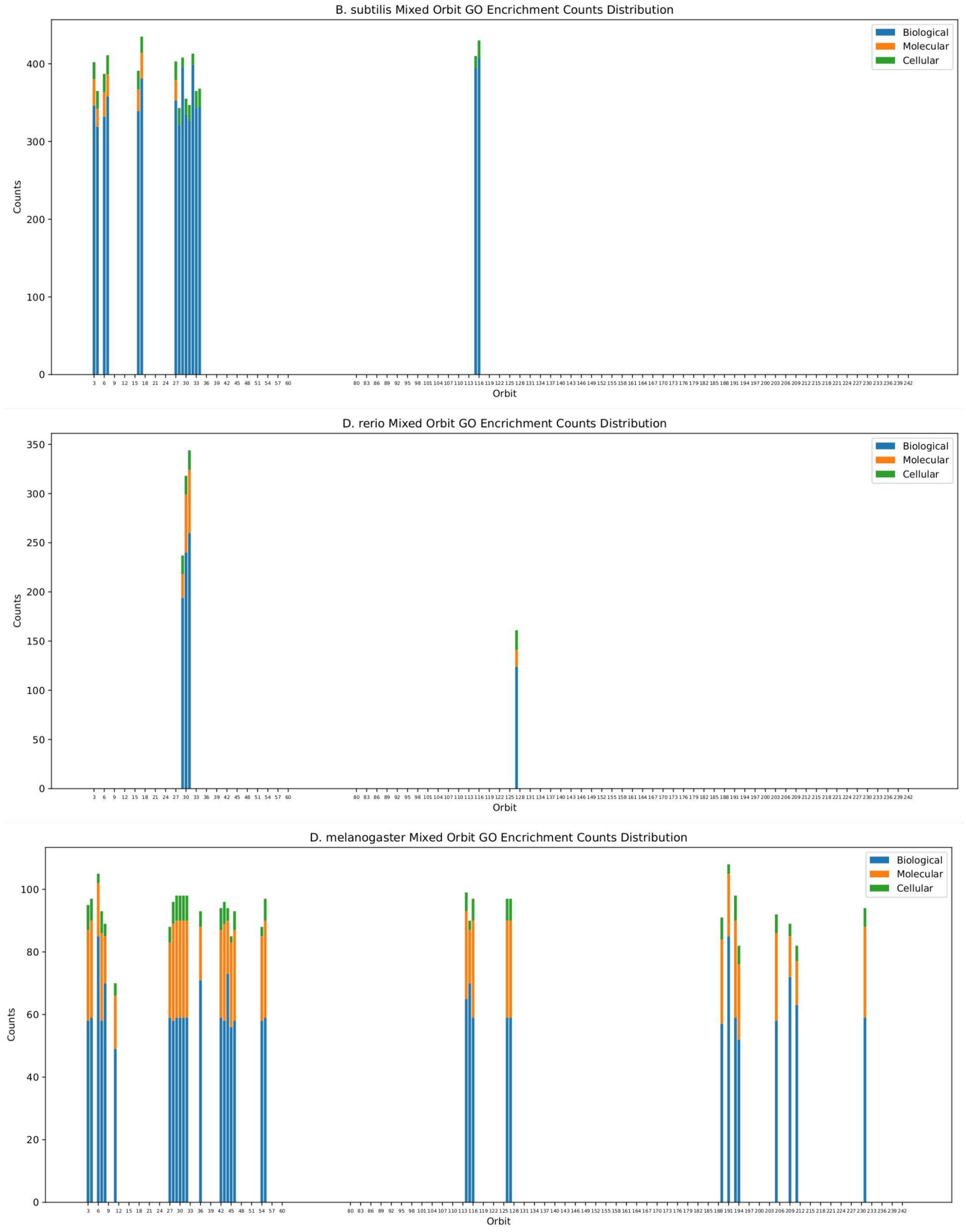

Figure S12: Mixed Orbit GO enrichment distribution across *B. subtilis*, *D. rerio*, and *D. melanogaster*, with stacked bar chart of GO enrichment counts for three main GO categories, Biological Process, Molecular Function, and Cellular Component

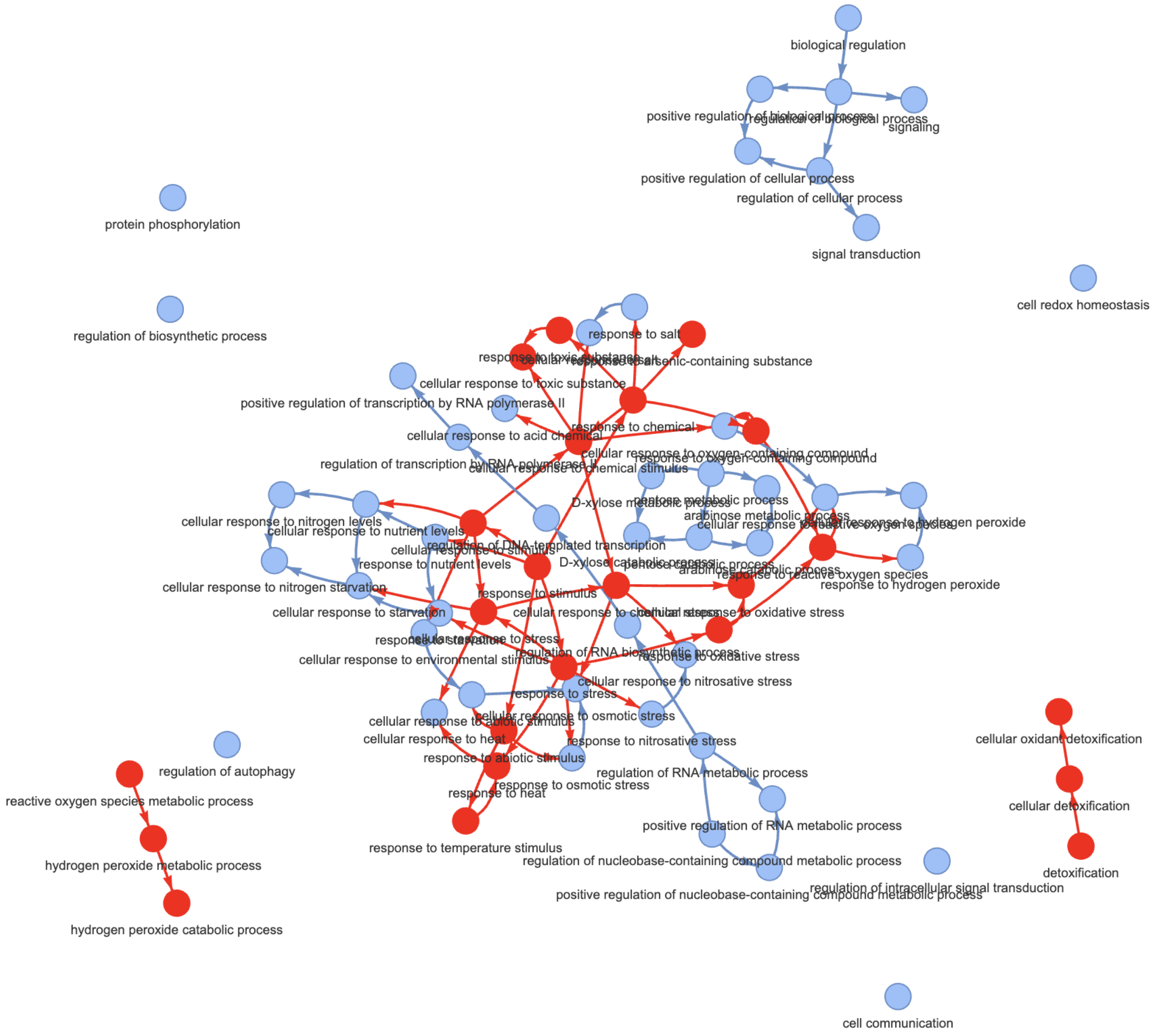

Figure S13: Graph visualization of GO term enrichment results for $O_{115}$ and $G_{45}$ for response to oxidative stress proteins. Red nodes indicate GO terms present in both results and blue nodes represent GO terms only found in $O_{115}$ results. Arrows are directed to show the hierarchy of GO terms.

#### References

[1] ConvertCSV.com - Convert CSV To JSON, XML, SQL, ...

[2] Uniprot: the universal protein knowledgebase in 2023. *Nucleic Acids Research*, 51(D1):D523–D531, 2023.

[19] Orsolya Liska, Balázs Bohár, András Hidas, Tamás Korcsmáros, Balázs Papp, Dávid Fazekas, and Eszter Ari. Tflink: an integrated gateway to access transcription factor–target gene interactions for multiple species. *Database*, 2022:baac083, 2022.

[20] Thilakam Murali, Svetlana Pacifico, Jingkai Yu, Stephen Guest, George G Roberts, and Russell L Finley. Droid 2011: a comprehensive, integrated resource for protein, transcription factor, rna and gene interactions for drosophila. *Nucleic acids research*, 39(suppl_1):D736–D743, 2011.

[21] Rose Oughtred, Jennifer Rust, Christie Chang, Bobby-Joe Breitkreutz, Chris Stark, Andrew Willems, Lorrie Boucher, Genie Leung, Nadine Kolas, Frederick Zhang, et al. The biogrid database: A comprehensive biomedical resource of curated protein, genetic, and chemical interactions. *Protein Science*, 30(1):187–200, 2021.

[22] Tiago Pedreira, Christoph Elfmann, and Jörg Stülke. The current state of subti wiki, the database for the model organism bacillus subtilis. *Nucleic Acids Research*, 50(D1):D875–D882, 2022.
